# Single-cell analysis of the epigenome and 3D chromatin architecture in the human retina

**DOI:** 10.1101/2024.12.28.630634

**Authors:** Ying Yuan, Pooja Biswas, Nathan R. Zemke, Kelsey Dang, Yue Wu, Matteo D’Antonio, Yang Xie, Qian Yang, Keyi Dong, Pik Ki Lau, Daofeng Li, Chad Seng, Weronika Bartosik, Justin Buchanan, Lin Lin, Ryan Lancione, Kangli Wang, Seoyeon Lee, Zane Gibbs, Joseph Ecker, Kelly Frazer, Ting Wang, Sebastian Preissl, Allen Wang, Radha Ayyagari, Bing Ren

## Abstract

Most genetic risk variants linked to ocular diseases are non-protein coding and presumably contribute to disease through dysregulation of gene expression, however, deeper understanding of their mechanisms of action has been impeded by an incomplete annotation of the transcriptional regulatory elements across different retinal cell types. To address this knowledge gap, we carried out single-cell multiomics assays to investigate gene expression, chromatin accessibility, DNA methylome and 3D chromatin architecture in human retina, macula, and retinal pigment epithelium (RPE)/choroid. We identified 420,824 unique candidate regulatory elements and characterized their chromatin states in 23 sub-classes of retinal cells. Comparative analysis of chromatin landscapes between human and mouse retina cells further revealed both evolutionarily conserved and divergent retinal gene-regulatory programs. Leveraging the rapid advancements in deep-learning techniques, we developed sequence-based predictors to interpret non-coding risk variants of retina diseases. Our study establishes retina-wide, single-cell transcriptome, epigenome, and 3D genome atlases, and provides a resource for studying the gene regulatory programs of the human retina and relevant diseases.

## Introduction

Retinal diseases, including age-related macular degeneration (AMD), diabetic retinopathy, glaucoma, and retinal vein occlusion, are major causes of vision loss in the United States, particularly affecting older adults and individuals with diabetes (*1*). AMD impacts 1.8 million Americans, while diabetic retinopathy affects 4.1 million (*1–3*). As these conditions are expected to increase with an aging population and the rising prevalence of diabetes, there is a critical need to develop effective early detection, prevention, and treatment strategies. The retina plays a dual role as a sensory interface and a processor of visual information (*4–6*). Genome-wide association studies (GWAS) have further revealed a strong genetic component to retinal diseases, identifying a large number of risk variants, the majority of which reside in noncoding regions of the genome (*7*). These noncoding variants are thought to modulate disease risks by altering the function of cis-regulatory elements (CREs) and gene expression patterns in retinal cell types (*8–10*). However, the lack of comprehensive annotation of CREs and their target genes across diverse retinal cell types remains a significant barrier to understanding the mechanisms by which these variants contribute to disease pathogenesis.

Recent advancements in single-cell genomic technologies have enabled detailed exploration of cellular heterogeneity within complex tissues, including the retina. Methodologies such as snRNA-seq/scRNA-seq and snATAC-seq/scATAC-seq (*11–16*), or 10x multiome single cell ATAC/RNA-seq (*7*, *17*) have been employed to investigate the complex regulatory mechanisms and disease pathology. Further, leveraging 3D genome data from HiChIP and Hi-C assays unraveled the cis-regulatory interactions (*18*, *19*). By integrating single-cell approaches with GWAS data, researchers are beginning to link noncoding variants to specific regulatory sequences in distinct retinal cell types, providing new insights into disease mechanisms. However, cell types of retinal tissues from younger donors remain unexplored(*20*, *21*). Additionally, knowledge of cell-type-specific methylation patterns, which can be influenced by environment (*22*), diet (*23*), and age (*24*), is still incomplete, hindering deeper understanding of the role of epigenetic processes in eye development and disease (*25*, *26*).

In this study, we comprehensively characterized the epigenome and 3D chromatin architecture of human retinal cell types using fresh post-mortem retinal tissues, collected within two hours of donor death, from three donors aged 20 to 40. We performed single-nucleus multiome (snATAC-seq/snRNA-seq) and single-nucleus methyl-3C sequencing (snm3C-seq) experiments (*29*, *30*), profiling gene expression, chromatin accessibility, DNA methylation, and chromatin conformation in over 58,000 retinal cells. Integrative multi-omic analysis of these datasets identified 420,824 candidate CREs (cCREs), revealing their cell-type-specific usage and potential target genes across 23 retinal cell subtypes from retina, macula and RPE/choroid tissues. Leveraging GWAS data, we identified cell types relevant to a spectrum of eye diseases, and determined likely causal SNPs for AMD and Macular telangiectasia (MacTel). Comparative analysis between human and mouse chromatin landscapes uncovered rapid turnover of gene regulatory elements during evolution. Additionally, we developed a deep neural network model to predict the regulatory functions of disease risk variants, and validated the predictions using CRISPR editing and hTERT-RPE1 cells. These data are publicly accessible and can be visualized via a custom built Web Portal (https://epigenome.wustl.edu/EyeEpigenome/) (*27*, *28*).

## Results

### A single-cell epigenome atlas of human retina

To create a comprehensive single-cell atlas of the human retinal epigenome, we performed single-nucleus 10x multiome (10x Genomics snRNA-seq and snATAC-seq) and snm3C-seq experiments on retina, macula, and retinal pigment epithelium (RPE)/choroid tissues from three phenotypically healthy donors (Fig. 1A). These methodologies allowed us to simultaneously profile transcriptomes alongside chromatin accessibility or DNA methylation with 3D genome organization in the same cells (*29*, *30*). In total, we analyzed 34,230 nuclei from the retina and macula using 10x multiome, with an additional 4,579 nuclei from the same tissues using snm3C-seq (Fig. 1B). For RPE/choroid tissues, 18,176 nuclei were profiled with 10x multiome, and 2,045 nuclei with snm3C-seq (Fig. 1C). The four modalities of Muller glia and RPE cell types are shown as examples (Fig. 1D).

**Fig. 1.**
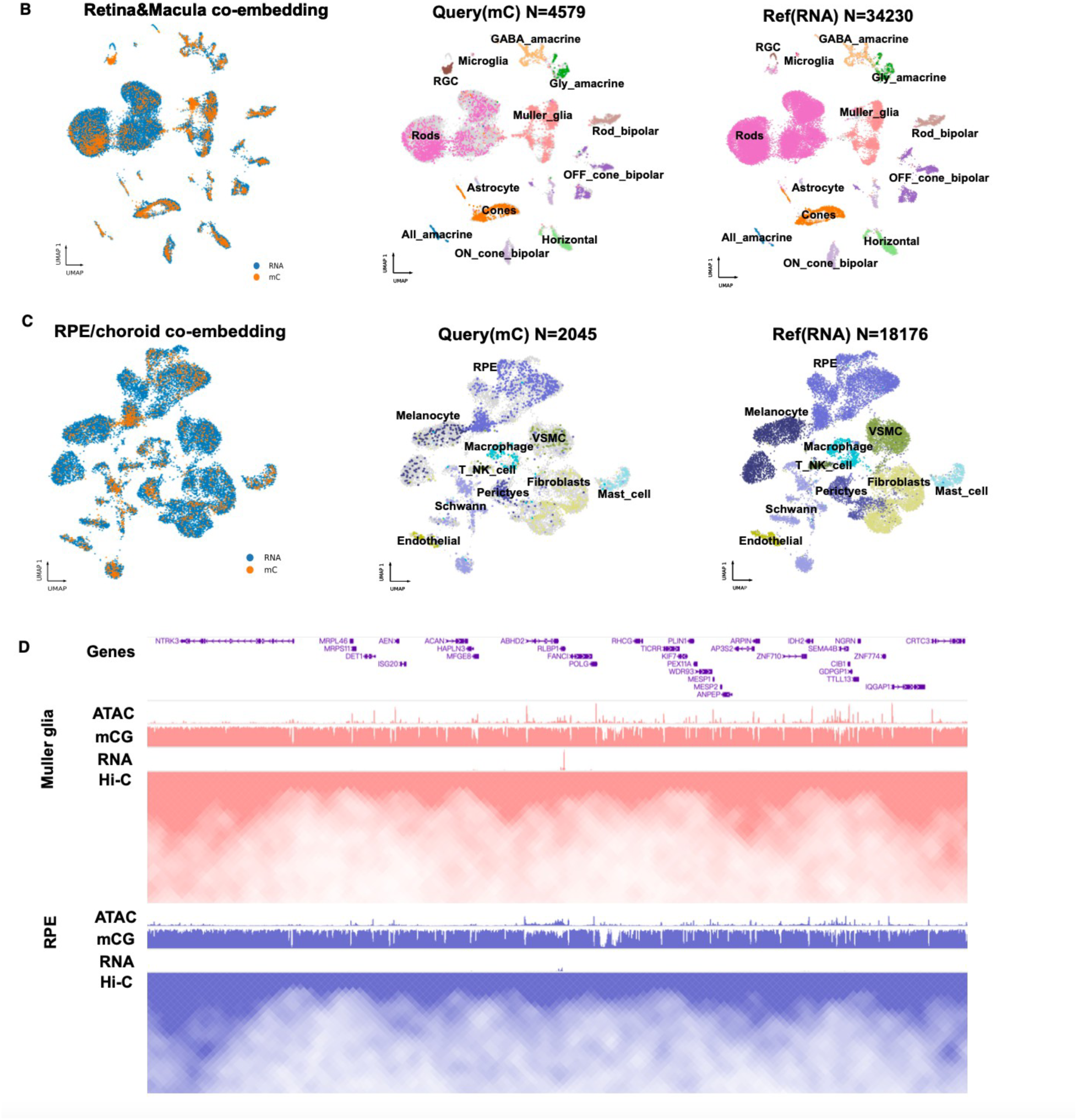
Single-cell multiomics analysis of the human retina. (**A**) Illustration of the human retina (left), specific cell type locations (middle), and experimental design. Tissues from three donors were used for 10x-multiome and snm3C-seq experiments (right). (**B**) Uniform manifold approximation and projection (UMAP) embeddings of 10x multiome RNA and snm3C-seq DNA methylation from human retina and macula. See method for details of clustering and integration of the two types of datasets(**C**) UMAP embeddings of 10x multiome and snm3C-seq data from human RPE. See methods for details of the integration and clustering of two types of datasets (**D**) Visualization of the pseudo bulk signals of gene expression, chromatin accessibility, DNA methylation and chromosome conformation using the WashU EyeEpigenome Browser. Muller glia and RPE cell types are shown as examples.

We conducted unsupervised clustering with the snRNA-seq component of the 10x multiome data from the retina and macula, resulting in 13 distinct cell clusters. We annotated the cell identity of each cluster according to known cell-type marker gene expression (*7*), including Rods, Cones, rod bipolar cells, OFF cone bipolar cells, ON cone bipolar cells, Muller glia, horizontal cells, GABA amacrine cells, Glycine amacrine cells, Retinal Ganglion Cells (RGC), All amacrine cells, astrocytes, and microglia (fig. S1, A, B).

For DNA methylome clustering (snm3C-seq), we annotated 13 cell types across 4,579 nuclei in the retina and macula tissues, guided by hypomethylation levels of marker genes and integration with the snRNA-seq dataset (methods). The RPE/Choroid dataset was grouped into 10 distinct cell types across 2,045 nuclei (Fig. 1B, C).

Next, we assigned cell-type identity to each cell cluster based on expression of known marker genes (Fig.2A, table S1). Across 30,293 detected genes, we identified 14,390 differentially expressed genes (Fig. 2B, table S2, methods). We also identified the top 50 marker genes per cell type using Seurat (*31*) (fig. S1C, methods). Gene Ontology (GO) enrichment analysis of these markers revealed expected functional terms, such as “detection of visible light” for Rods (Fig. 2C) and “visual perception” for Cones (Fig. 2D) (*4*).

**Fig. 2.**
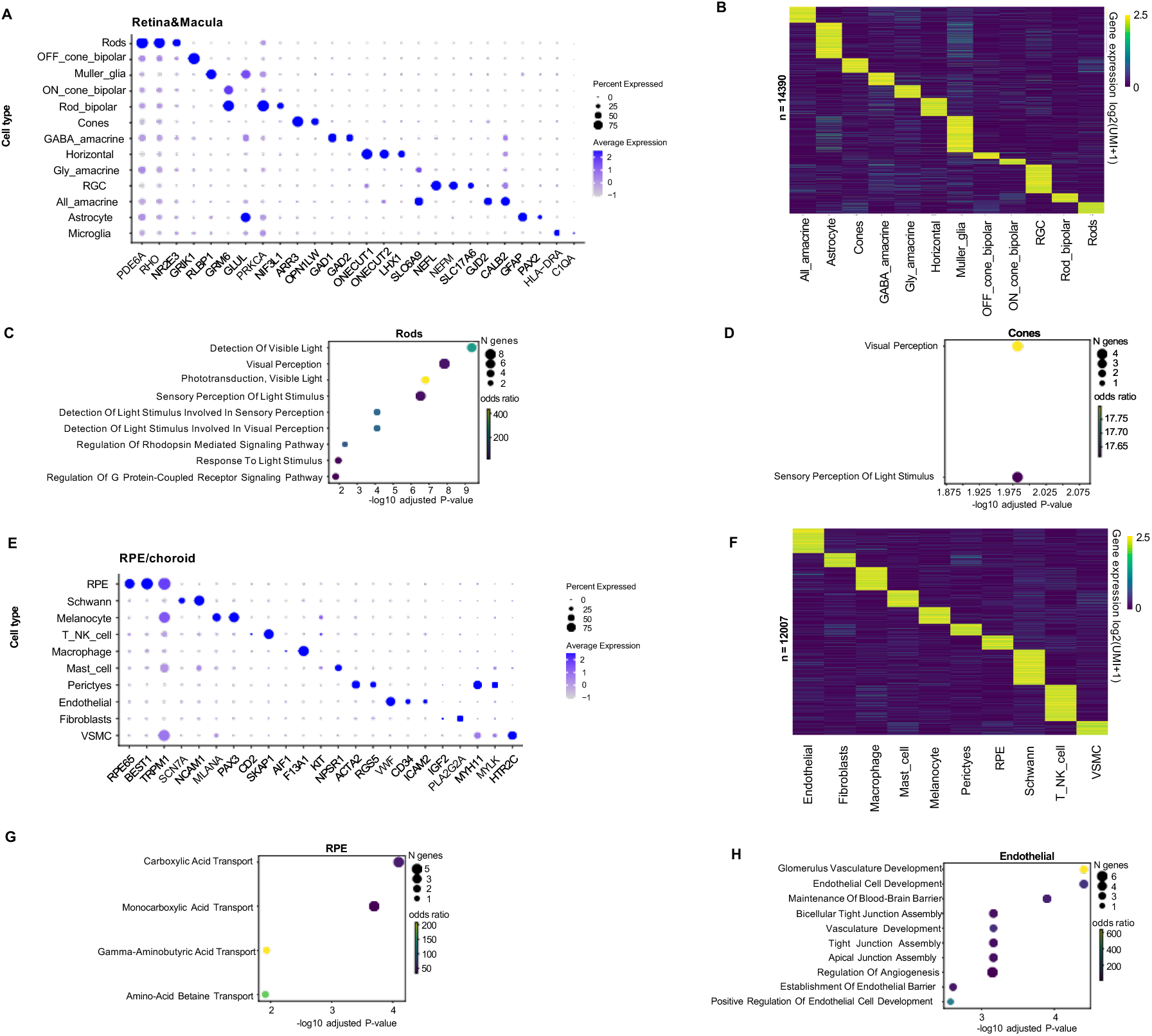
Transcriptional profiles from different cell types of the human retina and macula. (**A**) Dot plot visualizing the normalized RNA expression of selected marker genes in each cell type of retina and macula tissues. The color and size of each dot correspond to the average expression level and fraction of expressing cells. (**B**) Heatmap showing the expression of cell-type specific genes detected in retina and macula tissues. UMI, unique molecular identifier. Here the UMI is processed with the CPM method. The color bar shows the color gradient for the expression levels, with a score over 2.5 colored with 2.5. See method for details of the computation. (**C**) The significant GO terms for marker genes of Rods cells. **(D)**The significant GO terms for marker genes of Cones. **(E)** Dot plot visualizing the normalized RNA expression of selected marker genes in each cell type of RPE/choroid tissue. (**F**) Heatmap showing the expression of cell-type specific genes detected in RPE/choroid tissue. (**G**) Top significant GO analysis terms for marker genes of the RPE cell type. (**H**) Top significant GO analysis terms for marker genes of the Endothelial cell type.

In RPE/choroid tissue, we identified 10 distinct cell types (RPE, Schwann, Melanocyte, T/NK cell, Macrophage, Mast cell, Pericytes, Endothelial cells, Fibroblasts, and VSMC) based on snRNA-seq data and known marker genes (fig. S1D, Fig. 2E, table S3). Among the 30,037 genes profiled in the RNA assay, 12,007 were identified as differentially expressed (Fig. 2F, table S4, methods). The top 50 marker genes per cell type from RPE/choroid were determined using Seurat (*31*) (fig. S1E), with consistent expression across donors. GO analysis of these top 50 RPE cell marker genes revealed functional categories consistent with cell type identity, such as “gamma-aminobutyric acid transport” for RPE cells (Fig. 2G) and “glomerulus vasculature development” for Endothelial cells (Fig. 2H) (*4*).

Notably, identified marker genes *RLBP1* (*Retinaldehyde Binding Protein 1*) in Müller glia and *RPE65* in RPE are associated with the visual cycle (*32*) and human retinal disease (*33*), highlighting their homogeneous high expression cross different donors in Muller glia or RPE than other cell types (fig. S1, F, G, with additional data in fig. S7 and fig. S8).

### Identification and characterization of candidate *cis*-regulatory elements in retina cell types

To define gene-regulatory programs across distinct retina and RPE/choroid cell types, we identified candidate *cis*-regulatory elements (cCREs) by profiling open chromatin in 22 cell types (excluding microglia due to a low cell number). For each cell type, we aggregated snATAC-seq fragments and used MACS2 (*34*) to identify accessible chromatin regions. Previous research indicates that cluster size and read depth can impact MACS2 peak scores (*35*), with approximately 1,000 nuclei required to capture over 80% of accessible regions. Accordingly, we set a q-value of 0.05 for clusters with more than 1,000 nuclei and q-value of 0.1 for clusters with fewer nuclei. We iteratively merged the open chromatin regions identified from every cell type and kept the summits with the highest MACS2 peak score for overlapped regions.

Across retina and macula cells, we detected 10,408 to 198,843 open chromatin regions per cell type (500 bp span), with a combined total of 302,856 unique regions across 12 cell types. Of these, 113,996 regions showed significant cell-type specific chromatin accessibility (Fig. 3A, table S5, methods). In RPE/choroid tissues, we identified 12,929 to 142,274 open chromatin regions per cell type, with a union of 229,320 regions across 10 cell types. Among these, 67,299 displayed significant cell-type specificity (Fig.3B, table S6, methods). Notably, 111,352 accessible regions are shared between RPE/choroid and retina/macula tissues. Altogether, we identified 420,824 unique cCREs from retina, macula, and RPE/choroid tissues, most of which displayed highly cell type-specific patterns of accessibility (Fig. 3C-G).

**Fig. 3.**
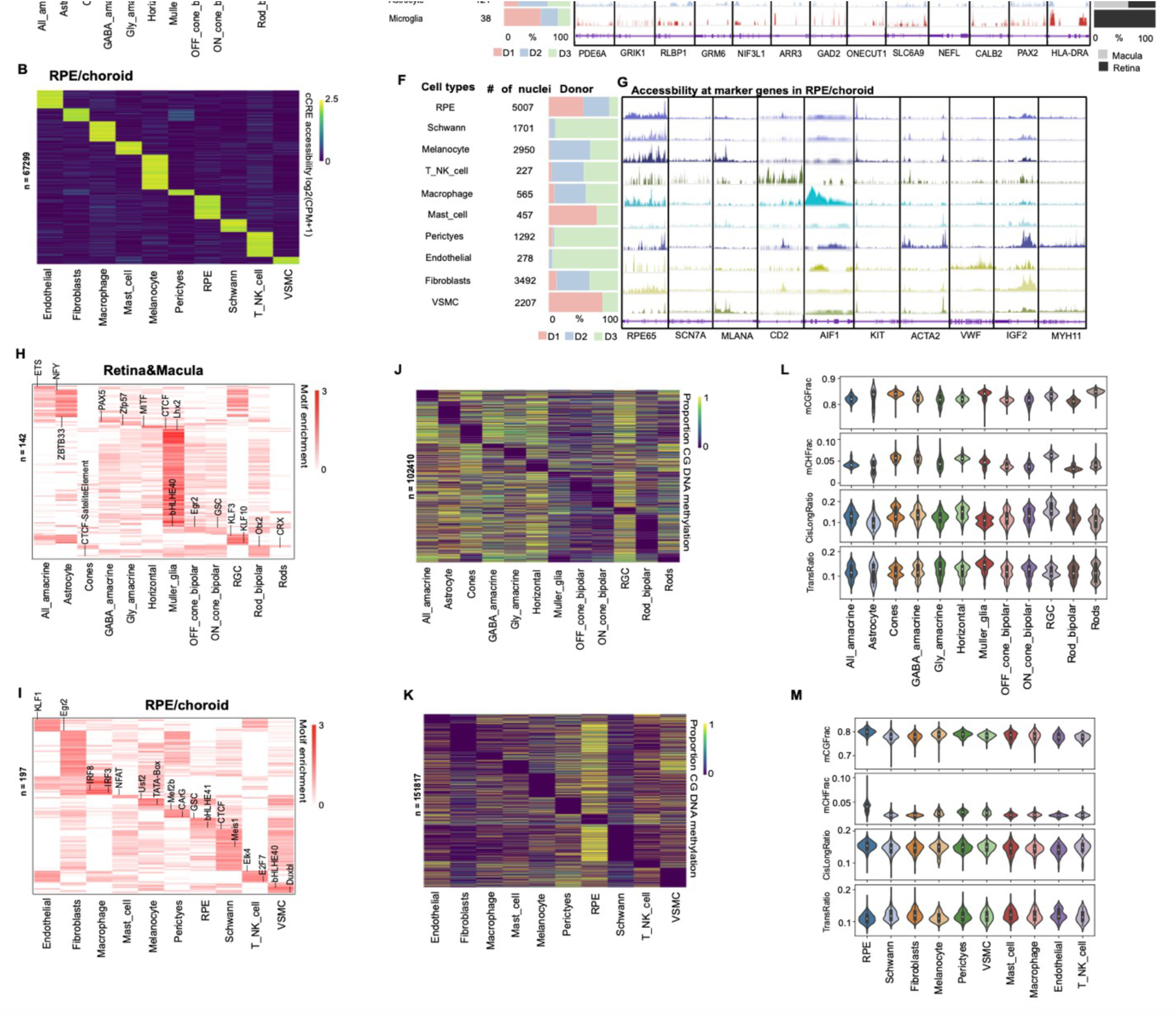
Identification and characterization of cCREs, TF motifs and DMRs across human retina cell types. (**A**)Heatmap showing chromatin accessibility of cell-type-specific cCREsof retina and macula tissues. CPM, counts per million. Each CRE is ordered by the cell type with the highest accessibility level. The color bar shows the color change for the accessibility change, a score over 2.5 is colored with 2.5. See method for details of the computation. (**B**) Heatmap showing chromatin accessibility of cell-type-specific cCREs of RPE/choroid tissue. (**C**) Stacked bar chart showing the contribution of each donor to each cell type in the retina/macular tissue. (**D**) Genome browser tracks chromatin accessibility profiles for each cell type of retina and macular tissues at selected marker gene loci. (**E**) Stacked bar chart representing the relative contribution of retina and macula regions to each cell type. (**F**) Stacked bar chart showing the contribution of each donor to the cell counts of each cell type of RPE/choroid tissue. (**G**) Genome browser tracks of chromatin accessibility profiles for each cell type at selected marker gene loci that were used for cell cluster annotation of RPE/choroid tissue. (**H** and **I**) Enrichment of TF motifs in cell-type-specific cCREs of retina and macula tissues (H) and RPE/choroid tissue (I). (**J** and **K**) Heatmap showing the DMRs of retina and macula tissues (J) and RPE/choroid tissue(K). Each DMR is ordered corresponding to the existing cCREs. The max value is 1. **(L** and **M)** Violin plot of mCG, mCH, CisLongRatio and TransRatio of each cell type in retina and macula tissues (L) and RPE/choroid tissue (M).

To explore the potential regulators of these cCREs, we performed motif enrichment analysis on the accessible chromatin regions for each cell type. In retina and macula tissues, 142 known transcription factor (TF) motifs were enriched within the cCREs across the 12 cell types, most of which displayed cell-type-specific enrichment patterns (Fig.3H, table S7, methods). In RPE/choroid tissue, 197 known motifs were enriched among cCREs across 10 cell types, most with cell-type-specific enrichment (Fig. 3I, table S8, methods). For retina and macula, certain TFs, such as Otx2, are known to play roles in both rod cells and rod bipolar cells (*36*). In RPE/choroid tissue, many of these TF motifs, such as Otx2 and CRX in RPE cells, have been previously implicated in cell-type-specific gene regulation (*37*).

This comprehensive list of candidate TF regulators provides a valuable resource for understanding gene-regulatory networks in human retinal and RPE cell types, offering insights for further studies into the regulatory elements that drive cell-type-specific functions in the retina and their implications for retinal diseases.

### DNA methylomes across the retina cell types

DNA methylation, or 5-methylcytosine (5mC), frequently present at cytosine-guanine dinucleotides (CpGs), is a key epigenetic modification involved in gene regulation and cell-type-specific functions within the retina (*38*). Differentially methylated regions (DMRs) across cell types are enriched at cCREs (*39*, *40*) and have been used for identifying cCREs. In addition to CpG methylation (mCG), methylation in non-CG (mCH, where H = A, C, or T) contexts is abundant in certain cell types in particular neurons and plays a crucial role in cell-type-specific gene regulation (*41*). The DNA methylation profile of the human retina can also aid in identifying genetic variations associated with specific retinal diseases (*26*).

Using the methylation modality of snm3C-seq, we generated a single-cell atlas of DNA methylation in the human eye, spanning cell types from the retina, macula, and RPE/choroid (fig. S1, H, I). This atlas provides cell-type-specific DNA methylation patterns, offering insights into the regulatory landscape of each retinal cell type. DMRs were identified using the ALLCools software (*38*), resulting in 102,410 unique CRE-associated DMRs across the 12 cell types in retina and macula tissues (Fig.3J). In RPE/choroid tissue, we identified 151,817 unique CRE-associated DMRs across 10 cell types (Fig.3K). These DMRs typically displayed an anti-correlated pattern with cCREs, reinforcing the cell-type specificity of DNA methylation in the retina. Consistent with previous findings (*42*), we observed that neuronal cell types (e.g., Rods, Cones, Horizontal, Rod bipolar, ON cone bipolar, OFF cone bipolar, All amacrine, GABA amacrine, Gly amacrine, and RGC) exhibited significantly higher levels of non-CG methylation (mCH) compared to non-neuronal cell types (e.g., Müller glia, Astrocytes, RPE, Melanocytes, T/NK cells, Mast cells, Pericytes, Fibroblasts, Endothelial cells, Schwann cells, VSMC, and Macrophages) (Fig. 3L, M).

### 3D genome architecture across retinal cell types

The three-dimensional (3D) organization of chromatin plays a critical role in gene regulation, with chromatin structures including active (A) and repressive (B) compartments, topologically associating domains (TADs), and chromatin loops influencing interactions between gene promoters and distal regulatory elements (*43*). These structures are also involved in essential nuclear processes, including DNA repair, homologous recombination, and replication (*42*, *44*). Eva D’haene et al. had explored the role of tissue-specific 3D genomic structures in establishing retinal disease gene expression patterns in the neural retina and RPE/choroid (*18*). Here, we sought to explore the cell-type-specific 3D genomic architecture of retinal cell types and reveal specific regulation mechanisms.

To investigate cell-type-specific genome folding, we first examined chromatin contact frequencies across different genomic distances, which is usually relevant to the cell cycle (*45*). In both retina and macula tissues, most cell types displayed an enrichment of chromatin contacts at mid-range (200 kb to 2 Mb) and long-range (20 Mb to 50 Mb) scales (Fig. 4A). However, the ratio of mid- to long-range contacts varied across cell types, indicating differences in chromatin organization (Fig.4B). Similarly, RPE/choroid cell types showed enrichment of contacts in both mid- and long-range categories, with distinct mid- to long-range ratios across cell types (Fig.4C, 4D). These contact profiles could provide insights into gene regulation in different cell types. We analyzed chromatin compartments and TADs in various retinal and RPE/choroid cell types, defining compartments at a 100-kb resolution (Fig.4E, 4G) and TAD boundaries at a 25-kb resolution. The boundary probability of a genomic bin, representing its frequency as a TAD boundary across cells, aligned closely with insulation scores derived from cell-type pseudo-bulk contact maps (Fig.4F, 4H). Notably, cell-type-specific compartment and TAD boundary patterns were evident, particularly around cell-type marker genes like *RLBP1* in Müller glia cells and *RPE65* in RPE cells. In regions approximately 2 Mb upstream of the transcription start site (TSS) and downstream of the transcription end site (TES) of these marker genes, we observed distinct domain boundary patterns across cell types, reflecting cell-type-specific chromatin organization (Fig.4F, 4H).

**Fig. 4.**
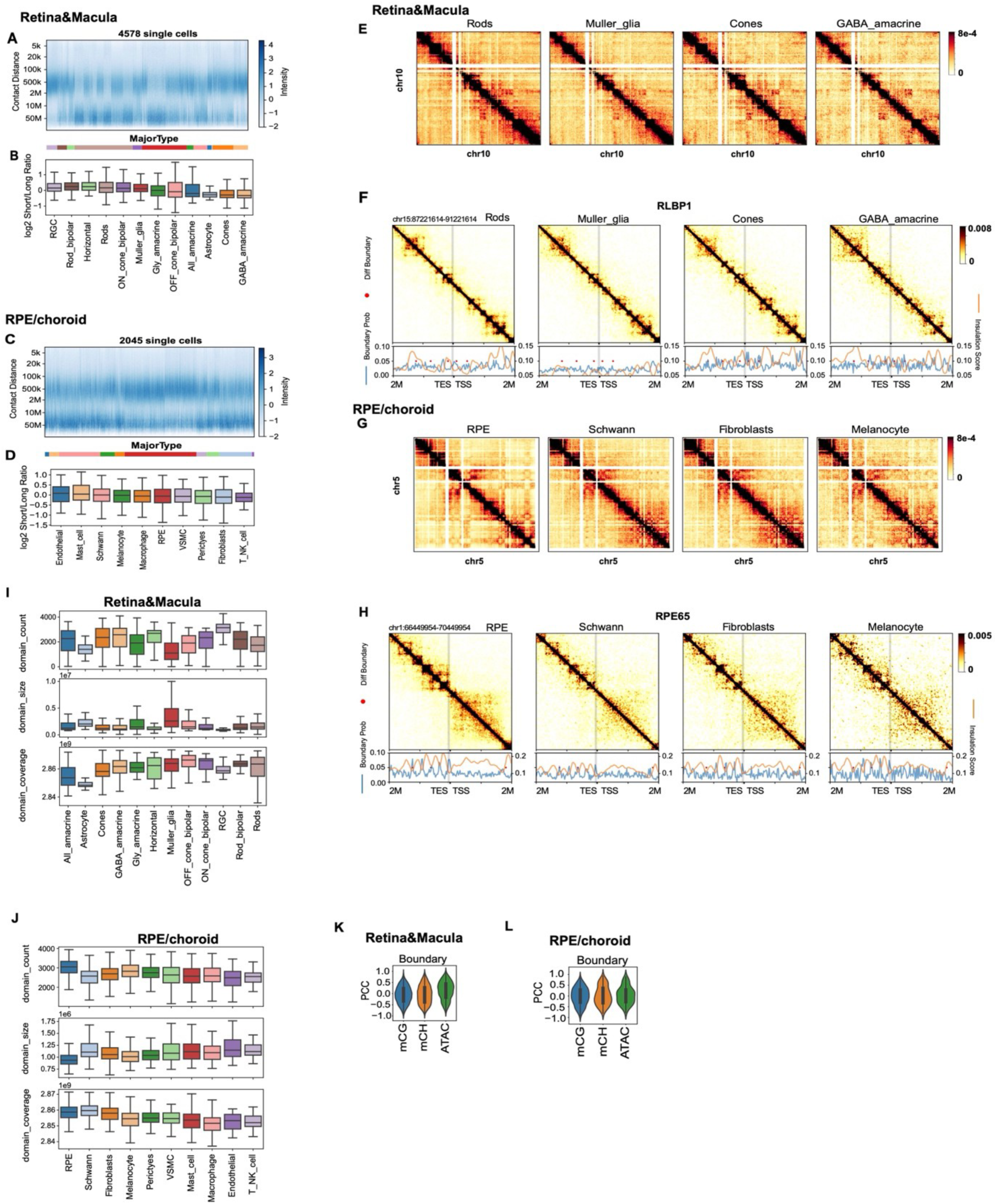
3D genome architecture across different retinal and choroidal cell types. (**A**) Frequency of chromatin contacts at different genomic distances in each single cell of retina and macula tissues. The color bar of intensity is Z-score normalized within each cell (column). The cells are grouped by cell type and then ordered by the median log2 short/long ratio over cells. The y axis is binned at log2 scale. (**B**) Box plots showing the distributions of Log2 short/long ratios of chromatin contacts in each cell type of retina and macula tissues, ordered the same as in (a). (**C**) Frequency of chromatin contacts at different genomic distances in each single cell of RPE/choroid tissue. (**D**) Box plots showing the distributions of Log2 short/long ratios of chromatin contacts in each cell type of RPE/choroid tissue, ordered the same as in (C). (**E**) Pseudobulkcontact maps of four cell types of retina and macula tissues. (**F**) Imputed contact matrices (heatmap), boundary probabilities (blue lines), insulation scores (orange lines) of four cell types of retina and macula tissues at RLBP1 locus (a marker of Muller glia). Differential boundaries were noted as red dots in line plots. (**G**) Pseudo bulk contact maps of four cell types of RPE/choroid tissue. (**H**) Imputed contact matrices (heatmap), boundary probabilities (blue lines), insulation scores (orange lines) of four cell types of RPE/choroid tissue at the RPE65 locus (a marker of RPE). (**I and J**) Box plots showing the domain coverage, size and count of cell types in retina and macula tissues (I) and RPE/choroid tissue (J). (**K** and **L**) Pearson correlation coefficient (PCC) between boundary probability and ATAC signals, mCG and mCH fractions of the bin(s) across all cell types for retina and macula tissues (K) and RPE/choroid tissue (L).

For retina and macula tissues, we identified 1,792 variable TAD boundaries across the 12 cell types (Fig.4I, fig. S2A). In RPE/choroid tissues, we found 216 variable TAD boundaries across the 10 cell types (Fig.4J, fig. S2B). These differences in TAD boundaries may reveal cell-type-specific regulatory landscapes, emphasizing the distinct 3D genome organization in each retinal cell type.

We next examined the relationship between TAD boundaries and other epigenetic modalities, such as open chromatin, mCG, and mCH methylation. In retina and macula cell types, both mCG and mCH methylation were generally anti-correlated with TAD boundary probabilities (Fig.4K). In contrast, open chromatin signals showed positive correlations with boundary probabilities, indicating that accessible chromatin DNA hypomethylation is often enriched at TAD boundaries.

Similarly, in RPE/choroid cell types, mCG and mCH remained anti-correlated with boundary probabilities, while open chromatin showed a weaker positive correlation (Fig.4L).

### Linking distal cCREs to target genes

To understand the transcriptional regulatory programs underlying cell-type-specific gene expression in the human retina, we used the activity-by-contact (ABC) method (*46*) to link distal cCREs to their potential target genes. By integrating chromatin accessibility and contact maps across retina cell types, we identified 302,856 distal cCREs linked to 32,766 potential target genes, resulting in 207,616 cCRE-gene pairs (table S9, fig. S1M and table S10, methods). Among these, 39,218 pairs with the highest ABC scores exhibited strong cell-type specificity, revealing distinct regulatory landscapes for each cell type (fig. S1L). In RPE/choroid cell types, we connected 229,320 distal cCREs to 32,815 target genes, resulting in a total of 197,458 cCRE-gene pairs, with 49,210 pairs showing clear cell-type-specific patterns based on high ABC scores (fig. S1O and table S11, fig. S1P and table S12, methods). Additionally, DMRs associated with these cCREs exhibited cell-type specificity, reinforcing the unique regulatory networks in each retinal and RPE/choroid cell type (fig. S1, N, Q).

Our findings provide a comprehensive map of cCRE-gene interactions in human retina and RPE cell types, highlighting the intricate regulatory networks that drive cell-specific gene expression. The varied correlation patterns between cCRE accessibility, target gene expression, and ABC scores suggest that multiple regulatory factors contribute to gene expression, underscoring the need for further investigation into the complex mechanisms underlying retinal cell-type-specific gene regulation.

### Single-cell epigenome analysis of mouse retina

Studying gene conservation across species provides insights into the evolutionary origins of key genes, identifies essential developmental pathways, and helps guide the selection of suitable animal models for studying human diseases (*47*). To explore the evolutionary dynamics of cCREs in human retinal cells and evaluate the suitability of mouse models for studying human retinal diseases, we performed snRNA-seq and snATAC-seq separately on retina tissues from four adult mice (two each at 2.5 and 5 months of age, with one male and one female per age group). This approach enabled us to profile the transcriptome and chromatin accessibility assays at single cell resolution separately.

Using unsupervised clustering based on single-cell RNA-seq data, we identified 13 distinct cell types in the mouse retina (fig. S2C, methods). Each cluster was annotated according to known cell-type markers in the retina (*7*, *47*). After removing doublets, the annotated cell types included Rods, OFF cone bipolar cells, Müller glia, ON cone bipolar cells, Rod bipolar cells, Cones, GABA amacrine cells, Horizontal cells, Gly amacrine cells, Retinal Ganglion Cells (RGC), All amacrine cells, Astrocytes, and Microglia. We observed cell-type-specific expression of known cell type marker genes (fig. S2D, table S13). Label transfer from RNA-seq data onto ATAC-seq clusters enabled chromatin accessibility profiling within the same 13 cell types across 27,359 nuclei (fig. S2E, methods).

In the mouse retina, we identified a union of 124,056 open chromatin regions across the 13 cell types. Among these, 53,899 were significant cell type-specific cCREs (fig. S2F, methods) and likely regulate cell-specific gene expression within the retina. Additionally, we detected 23,295 expressed genes in the RNA-seq assay, of which 12,274 displayed significant cell type specific expression (fig. S2G, methods). This epigenomic and transcriptomic atlas enriches our understanding of cell-specific functions within the mouse retina.

To identify potential transcriptional regulators, we performed motif analysis on accessible regions across the 13 retinal cell types. A total of 168 known TF motifs were enriched among the cCREs from different cell types, most of which displayed cell-type-specific patterns (fig. S2H, table S14). Notable cell-type-specific motifs included ZNF143|STAF in All-amacrine cells; Sp1 in Astrocytes; CTCF in Cones; Tgif2 in GABA amacrine cells; Usf2 in Gly amacrine cells; ELF5 in Müller glia; Ronin in OFF cone bipolar cells; YY1 in ON cone bipolar cells; RORgt in RGC; Elk4 in Rod bipolar cells; and BORIS in Rods. These enriched motifs suggest regulatory roles for these TFs in specific retinal cell types, supporting distinct gene regulatory programs essential for retinal function.

### Comparative analysis of gene expression programs in human and mouse retinal cell types

To explore the conservation of gene-regulatory landscapes between human and mouse retinas, we performed a comparative analysis of chromatin states and gene expression in retinal cell types. Given that mouse eyes lack macula tissue (*48*), our analysis focused on merged data from human retina and macula tissues and mouse retina tissue. We profiled 34,230 cells from human retina and macula RNA-seq data and 27,359 cells from mouse retina RNA-seq data, using joint clustering based on gene activity scores to align cell types across species (Fig. 5A-C, fig. S1J-K, methods).

**Fig. 5.**
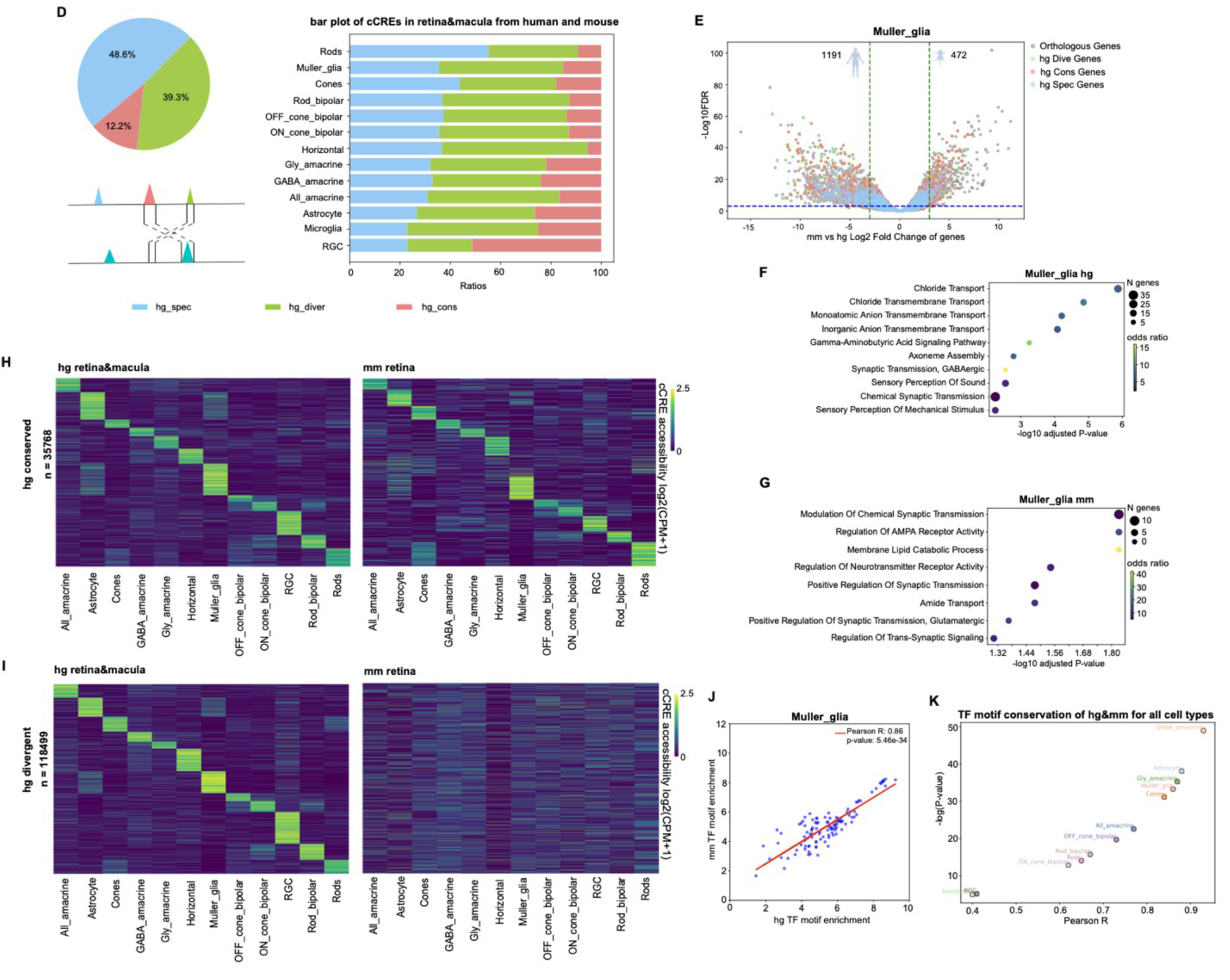
Comparative analyses of chromatin accessibility and gene expression between human and mouse retinal cell types. (**A**) UMAP co-embedding of single nuclei RNA-seq data, with each cell colored by species. (**B**) UMAP embedding of 10x multiome RNA-seq data of human retina and macula tissues, annotated by cell type. (**C**) UMAP embedding of single-cell RNA-seq of mouse retina tissue, annotated by cell type. (**D**) Comparison of human and mouse retinal cCREs. Left, pie chart showing fraction of three categories of cCREs, including human specific, human divergent, and human conserved cCREs. The human conserved cCREs are those both with DNA sequences conserved across species and in open chromatin in orthologous regions. The human divergent cCREs are sequences conserved but the orthologous regions have not been identified as open chromatin regions in mouse retina. The human specific cCREs are those without orthologous sequences in the mouse genome. Right, bar plot showing three categories of cCREs in corresponding cell types from human and mouse. (**E**) The differential gene expression between human and mouse in Muller glia. Gray dots represent all the orthologous genes across human and mouse in this cell type; green dots represent the genes paired with human divergent cCREs; red dots represent the genes paired with human conserved cCREs; blue dots represent the genes paired with human specific cCREs. (**F**) Top significant GO terms for significantly differentially expressed genes in human Muller glia. (**G**) Top significant GO terms for significant differentially expressed genes in mouse Muller glia. **(H)** Heatmap showing chromatin accessibility of putative human conserved enhancers of retina and macula tissues for human and mouse. **(I)** Heatmap showing chromatin accessibility of putative human divergent enhancers of retina and macula tissues for human and mouse. **(J)** Dot plot of conserved TF motif enrichment of human and mouse in Muller glia. (**K**) Scatter plot of TF motif conservation of human and mouse for all retinal cell types. X axis represents Pearson R, y axis represents -log(P-value).

We next investigated the conservation of chromatin accessibility at cCREs between corresponding cell types in the human and mouse retina. For 51.44% of human cCREs, we identified mouse genome sequences with high similarity (defined as >50% of bases lifted over to the human genome) (Fig.5D). Among these conserved sequences, 12.18% also showed chromatin accessibility in at least one cell type from the mouse retina. We termed these cCREs with both DNA sequence similarity and chromatin accessibility as "chromatin accessibility conserved cCREs." The remaining 39.26% of human cCREs showed sequence similarity without conserved chromatin accessibility, which we termed "human divergent cCREs." Additionally, 48.56% of human cCREs lacked orthologous sequences in the mouse genome and were classified as "human-specific cCREs," although they may be conserved in other primates or mammals (Fig.5D, left). This general pattern aligns with previous reports on chromatin conservation across species (*49*).

Breaking down these categories by cell type, we observed consistent proportions of conserved, divergent, and human-specific cCREs across different retinal cell types (Fig. 5D, right). This cell-type-specific breakdown provides insight into the evolutionary dynamics of chromatin accessibility in the retina.

To examine species-specific and conserved gene expression programs, we performed differential expression and chromatin accessibility analyses using edgeR (*75*) for each cell type. In Müller glia, we observed notable differences in RNA expression levels, with 1,191 genes showing higher expression in human cells and 472 genes showing higher expression in mouse cells (Fig.5E). Genes paired with human-specific cCREs were primarily found in regions with high expression in humans or nonsignificant regions, while genes paired with conserved cCREs spanned all expression regions (Fig.5E). Human divergent cCREs were mostly associated with genes exhibiting high expression in humans, although some orthologous genes with mouse divergent cCREs showed elevated expression in mice, potentially reflecting cell-type-specific functions in the mouse retina.

GO analysis of differentially expressed genes revealed distinct functional profiles, with genes highly expressed in humans (Fig.5F) and in mice (Fig.5G) enriched for specific processes. This functional divergence underscores potential species-specific roles in retinal biology, even among orthologous cell types. For example, compared with mice, highly expressed genes of human Muller glia cell type have functions relevant to synaptic transmission and GABAergic (Fig.5F), but murine Muller glia have functions relevant to membrane lipid catabolic process (Fig.5G). Additional cell type comparisons and their specific gene functions are provided in fig. S3.

For the 35,768 human-conserved cCRE regions, heatmaps revealed consistent cell-type-specific accessibility patterns across species (Fig.5H). In contrast, the 118,499 human-divergent cCRE regions displayed a clear cell-type-specific accessibility pattern in human retina cell types, while corresponding mouse sequences showed minimal chromatin accessibility (Fig.5I). This divergence in chromatin accessibility highlights epigenetic differences between species that may reflect adaptation to unique visual demands.

To investigate conservation at the level of TF motifs, we selected conserved TF motifs for each cell type and calculated the PCC of motif enrichment scores. For example, in Müller glia, we observed a Pearson correlation coefficient (R) of 0.86, indicating strong TF motif conservation between species (Fig. 5J). Similar analyses for other cell types (fig. S9A-K) demonstrated varying degrees of conservation. Overall, cell types with higher Pearson correlation coefficients tend to have lower p-values. For example, Horizontal and RGC cell types exhibit the lowest Pearson correlation coefficients and the highest p-values, suggesting a lower conservation of TF motifs between human and mouse in these cell types. In contrast, GABA amacrine cells show the highest conservation, as indicated by their higher Pearson correlation coefficient and lower p-value (Fig.5K). The conservation of TF motifs across species not only supports the use of mouse models to explore mechanisms underlying human retinal pathology but also strengthens their relevance in translational research.

### Epigenome maps facilitate the interpretation of non-coding risk variants

Mapping non-coding risk variants to specific cell types provides critical insights into the regulatory mechanisms underlying ocular diseases. By integrating identified cCREs and their associated transcription factor motifs, we can better understand chromatin accessibility across retinal cell types and assess the functional roles of non-coding regions linked to disease, particularly those identified by genome-wide association studies (GWAS).

GWAS have revealed genetic variants associated with numerous ocular diseases and traits, yet most of these variants are located in non-coding regions of the genome (*50*). Previous studies have shown that non-coding risk variants often overlap with cCREs active in disease-relevant cell types (*51*). Leveraging the newly annotated, cell-type-specific human retina cCREs, we first aimed to predict cell types associated with various eye diseases. Using linkage disequilibrium score regression (LDSC), we assessed whether genetic heritability of DNA variants linked to retinal diseases is significantly enriched within cCREs active in specific retinal cell types. This analysis revealed significant associations between seven eye diseases and cell-type-specific open chromatin profiles in the retina and macula tissues, and five in RPE tissue (Fig. 6A, 6B).

**Fig. 6.**
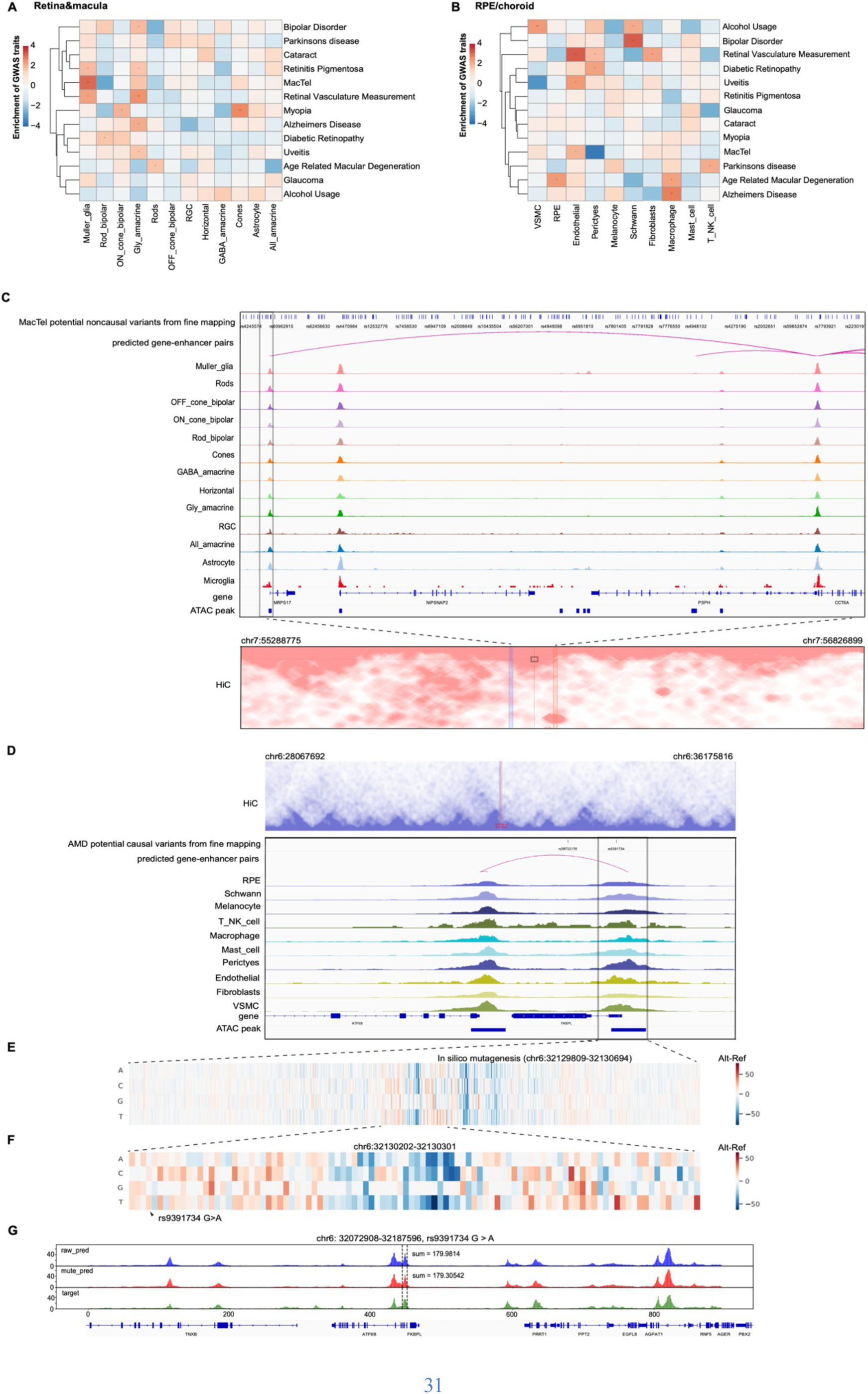
Interpreting noncoding risk variants of ocular disorders and traits. (**A** and **B**) Heatmap showing enrichment of risk variants associated with ocular disorders and traits in human cell-type-resolved cCREs of retina and macula tissues (A) and RPE tissue (B). LDSC analysis was performed using GWAS summary statistics of each disorder or trait. P values were corrected using the Benjamini-Hochberg procedure for multiple tests. FDRs of LDSC coefficients are shown. *FDR < 0.05; **FDR < 0.01; ***FDR < 0.001. (**C**) Genome browser tracks (GRCh38) display chromatin accessibility profiles from snATAC-seq and contact score map of a locus. Arcs represent the predicted pairs of enhancer and gene from ABC model in Muller glia cell type. (**D)** Genome browser tracks (GRCh38) display chromatin accessibility profiles from snATAC-seq and contact. Arcs represent the predicted pairs of enhancer and gene from ABC model in RPE cell type. (**E**) Chromatin influenced the prediction of accessibility predicted after *in silico* nucleotide mutagenesis using deep neural network models within region chr6: 32129809-32130694 (0-based) of RPE cell type. Red color represents increased accessibility predicted at the altered sequences, while blue color represents lower accessibility on altered sequences. (**F**) Zoom in of the region chr6: 32130202-32130301 of RPE cell type. Lower accessibility was predicted on a *FKBPL* enhancer with risk variant rs9391734 G>A. (**G**) Chromatin accessibility at *FKBPL* enhancer loci predicted in human RPE cell type. Green color represents the raw target track, blue represents predicted track, red represents predicted track after mutation.

Specifically, LDSC analysis using human-specific regulatory elements showed an association between MacTel and Müller glia cells (Fig.6A), suggesting that MacTel-related risk variants might reside within human-specific regulatory elements that drive gene regulation in Müller glia (*7*, *52*). Similarly, an association between age-related macular degeneration (AMD) and RPE cell types was observed (Fig.6B), indicating that AMD-related risk variants could influence gene regulatory programs specific to the human RPE (*53*).

To refine our understanding of disease-associated variants within cCREs, we conducted fine mapping for MacTel and AMD traits to prioritize lists of potential causal SNPs. For MacTel, we identified nine loci on chromosomes 1, 2, 3, 5, 7, 9, and 10, consistent with previous reports (*52*). Detailed visualization of these loci (fig. S6, K-S) revealed 118 likely causal SNPs. For AMD, we identified ten loci on chromosomes 1, 6, 8, 10, 16, and 19, also consistent with previous findings (*53*). Fine mapping with *susieR* (*54*) yielded a total of 88 likely causal SNPs across these AMD loci (fig. S6, A-J).

Given the strong association between Müller glia and MacTel in humans (Fig.6A), we hypothesized a similar relationship in the mouse retina. Surprisingly, no overlap was found between potential causal SNPs for MacTel and conserved human-mouse peaks. This lack of conservation indicates that, while Müller glia cells are present in both species, the causal gene regulatory variants linked to MacTel are probably human-specific. These findings underscore the necessity of using human or primate models to investigate the mechanisms underlying this disorder.

To further characterize SNP-target gene relationships, we examined overlaps between potential causal SNPs identified through fine mapping and peak regions from the ABC model, supported by Hi-C data. For MacTel, only one causal SNP directly overlapped with Müller glia peak regions (Fig.6C). For AMD, three RPE peaks overlapped with AMD-associated causal SNPs, including rs9391734, which overlapped with the peak region at chr6: 32,130,132-32,130,631 in RPE cells, paired with the target gene *ATF6B/FKBPL* (Fig.6D). Hi-C analysis further refined target gene predictions, demonstrating how the combination of single-cell multiomics with chromatin interaction data enhances our ability to interpret non-coding variants in ocular disease.

### Deep neural network predicts the effects of risk variants

To investigate how risk variants affect the function of regulatory elements, we adopted Basenji (*55*) to train a deep neural network (DNN) to predict chromatin accessibility from DNA sequences. We trained the DNN model on normalized pseudo-bulk ATAC-seq profiles from human RPE and melanocyte, selecting the model with the highest average PCC on validation dataset for each cell type. Specifically, our best model achieved PCC values of 0.8331 for the RPE dataset and 0.8353 for the melanocyte dataset (fig. S10, A, B, methods). We conducted *in silico* mutagenesis on RPE-specific cCREs associated with potential AMD-risk variants to assess their effects on chromatin accessibility across different scenarios. In the first case (Fig. 6D-G), we showed that the potential causal variant rs9391734, located within a peak region, slightly reduced the predicted accessibility of the *ATF6B/FKBPL* promoter, with high prediction accuracy (PearsonR = 0.915). In the second case (fig. S4, A-D), two potential variants (rs943079 and rs943080) within an enhancer, but without paired target genes, were predicted to either increase or decrease accessibility. In the third case (fig. S4, E-H), two potential variants (rs2672600 and rs3750847) located near but outside a peak region were found to have subtle regulatory effects on accessibility, with proximity to the peak summit correlating with their impact. In the fourth case (fig. S5, A-G), we observed that the same mutation within an enhancer could have different effects on accessibility across cell types, as demonstrated by divergent impacts in RPE and melanocyte cells. Additionally, deletion of enhancer sequences disrupted accessibility and altered neighboring enhancer activity, likely due to motif disruption (fig. S5H). These findings highlight the nuanced regulatory effects of both causal and noncausal variants on chromatin states across contexts.

To validate the deep neural network’s predictions for the effects of non-coding variants, we used CRISPR editing to genetically engineer the hTERT-RPE1 cell line. *TMEM216* is known to be associated with retinal degeneration and syndromes such as Joubert and Meckel (*56–60*). It is a component of cilia, specifically localized between the basal body and ciliary axoneme (*61*, *62*). Firstly using the model trained on the pseudo bulk chromatin accessibility profile of RPE cell type, we performed *in silico* mutageneisis, and predicted a decrease in chromatin accessibility following the c.-69G>A mutation (Fig. 7A,B). To evaluate the performance of our deep learning model, we used CRISPR editing to engineer two RPE1 cell lines harboring *TMEM216* c.-69G>A variant in homozygous state (D4 and F6), and heterozygous state (G1), and performed bulk ATAC-seq (Fig.7C). In agreement with the prediction, chromatin accessibility at this site was reduced in both homozygous cell lines D4, F6, and also in the heterozygous G1 cell line compared to the wild-type hTERT-RPE1 cells (Fig.7C). These findings are consistent with reduced levels of *TMEM216* expression reported in cells with c.-69G>A change (*56*). This result supports the utility of our deep learning model in predicting functional consequences of non-coding risk variants.

**Fig. 7.**
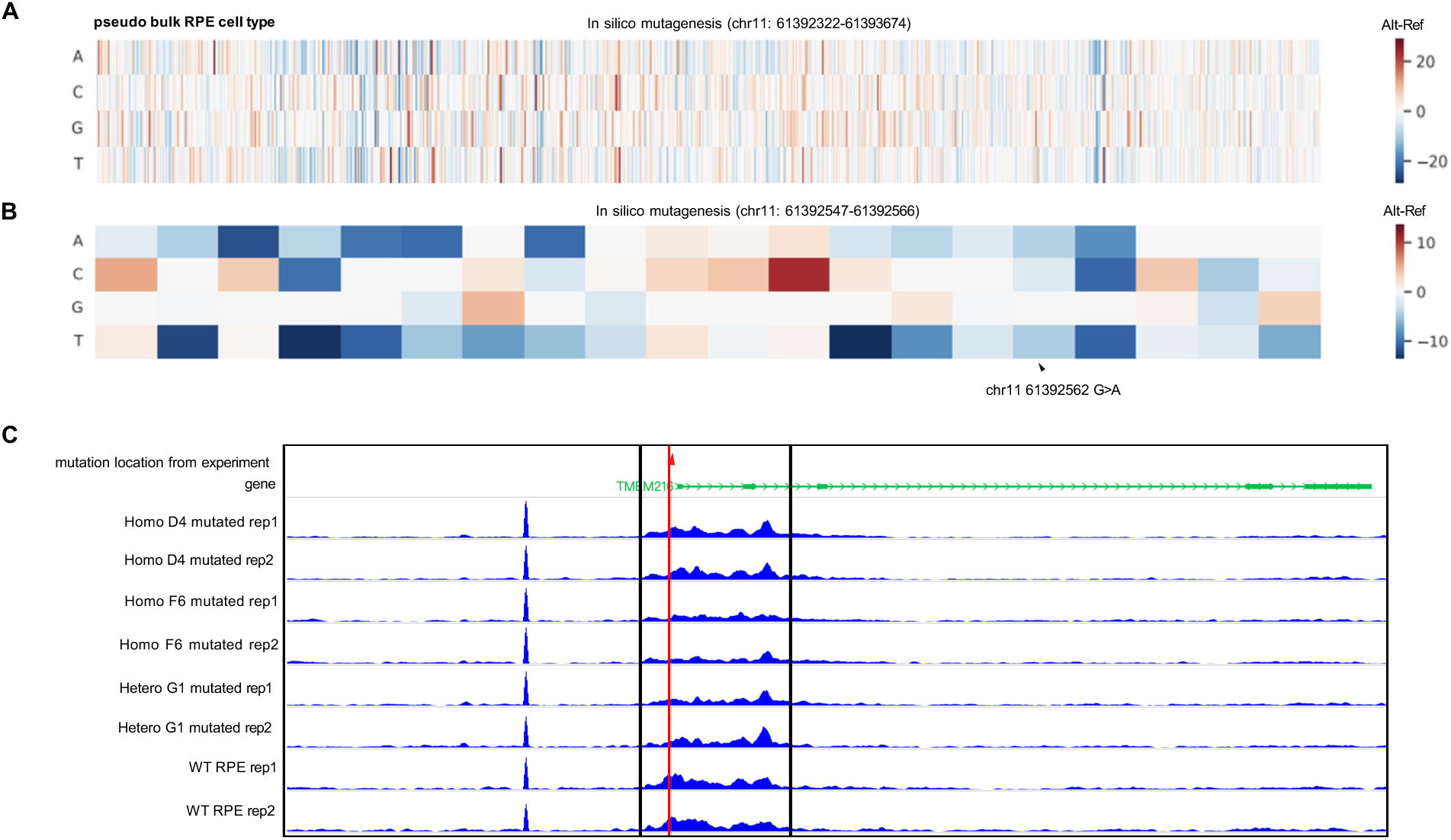
CRISPR editing experiments validate a prediction of the deep neural network model. (**A**) Predicted chromatin accessibility after *in silico* mutagenesis within region chr11: 61392322-61393674 (0-based) of pseudo bulk RPE cell type. (**B**) Zoom in *in silico* nucleotide mutagenesis within region chr11: 61392547-61392566. Lower accessibility was predicted on *TMEM216* enhancer with risk loci chr11 61392562 G>A. (**C**) Genome browser tracks (GRCh38) display chromatin accessibility profiles of CRISPR edited RPE cell lines. Two replicates were shown for each cell line.

### Discussions

Analysis of the transcriptome, epigenome and 3D genome features of 23 cell types from human retina tissues enhances our knowledge of gene regulatory programs in the human retina. Compared with other recent works, our study introduces several unique features:

First, by utilizing freshly collected retina tissues from young donors aged around 20–40 years, our study minimizes postmortem artifacts.

Second, we incorporated three tissues—retina, macula, and RPE/choroid in our analysis. The inclusion of RPE/choroid, rarely in previous studies, provides a more holistic view of retinal biology and its regulatory mechanisms. Leveraging four molecular modalities—gene expression, chromatin accessibility, DNA methylation, and chromatin conformation across 23 cell types, we report a resource for studying the cell type specific gene regulation of human retinal diseases analysis.

Third, we also provided detailed, cell-type-specific data, including values for CREs, genes, transcription factor motifs, and CRE-gene pairs, all presented in comprehensive tables. These resources are further enriched with cross-species information, offering a unique comparative perspective. Additionally, our extensive list of candidate transcription factors and their motifs, particularly those enriched in rare cell populations such as those in the RPE, lays a foundation for reconstructing detailed gene-regulatory networks and conducting targeted studies on retinal cell function and disease mechanisms. These curated datasets could facilitate research in retinal biology.

Fourth, our study represents the first report of cell-type-specific DNA single cell methylation data in the human retina, providing a valuable resource for linking DMRs with cCRE-gene pairs and disease-related loci. Future studies incorporating age-diverse and patient-derived samples could build on this work to reveal further insights into the role of cell-type-specific single cell DNA methylation in eye aging and disease.

Fifth, our deep cross-species comparisons highlight conserved enhancers and transcriptional programs, providing a framework to assess the utility of mouse models for human retina diseases.

Sixth, by integrating GWAS data, deep neural network predictions, and CRISPR validation, our analysis uncovers the genetic underpinnings of several eye diseases, highlighting the role of conserved and human-specific noncoding regulatory elements in polygenic traits. Using deep neural networks, we captured the gene-regulatory code and interpreted the effects of risk variants associated with complex traits and diseases. *In silico* mutagenesis enabled the identification of "high-effect" SNPs, offering insights into causal variants across various cell types. This integrative approach bridges computational predictions and experimental validation, facilitating the functional annotation of noncoding disease risk variants and advancing precision medicine for ocular diseases.

Finally, our study introduces a user-friendly web portal that enables researchers to explore the single cell multiomic data from 23 cell types across three eye tissues. This portal, enriched with cell type-specific information on cCREs, transcription factors, and 3D genome contacts, ensures accessibility and utility for a broad range of research applications.

Our study is limited by the small sample size, which includes only three individuals of varying ages. This limitation may impact the generalizability of our findings across broader demographic groups, including diverse ethnicities, genders, and age ranges. Expanding sample diversity and size in future studies will be crucial for refining our understanding of chromatin landscape variability across populations. Additionally, integrating single-cell multiomics with spatial transcriptomics will aid in identifying rare cell types and elucidating complex gene-regulatory networks, further enhancing our ability to pinpoint mechanisms through which genetic variants influence disease phenotypes.

In conclusion, our integrated approach advances the understanding of the genetic architecture of retinal diseases, demonstrating the potential of cutting-edge technologies to unravel complex gene regulation in polygenic human traits. As these findings are validated and expanded, the potential for significant clinical impact grows, promising to transform patient care through more personalized, genetically informed therapies. Our work also provides a strong foundation for future research on human retinal aging and the identification of retinal disease-associated variants.

## Acknowledgments

We thank Zhaoning Wang for bulk-ATAC seq experiment discussion and guidance; thank Jingtian Zhou for discussion; We would like to thank the families of donors and San Diego Eye Bank for providing the eye globes within 2 hours of enucleation.

## Author contributions

B.R. and R.A. supervised the study. Y.Y., P.B., Q.Y., A.W. generated the human 10x multiom data. Y.Y., N.R.Z.,P.B.,K.Y.D.,P.K.L.,W.B. generated the human snm3C-seq data. Y.Y., P.B., K.D.,J.B.,L.L., R.L.,A.W.,S.P.. generated the mouse 10x scRNA and scATAC data. P.B., Y.Y. and Z.G. prepared cell lines, Y.Y. generated bulk ATAC-seq data. Y.Y., N.R.Z., K.D., Y.W., M.D.A., Y.X., K.W.,S.L. analyzed the data. Y.Y., N.R.Z., P.B., R.A. and B.R. interpreted the data. D.L., C.S. and Y.Y., T.W. built the web portal. Y.Y. and B.R. wrote the manuscript. All authors edited and approved the final manuscript.

## Data and materials availability

Data produced in this study are available at the NCBI GEO under accession number GSE277326 (human eye 10x multiome), GSE277361 (human eye sn-m3C-seq), GSE276851 (human RPE cell lines bulk ATAC-seq), GSE276864 (mouse retina snRNA-seq), GSE276923 (mouse retina snATAC-seq). All the datasets are also available at the 4DN portal (https://data.4dnucleome.org/ren-lab-single-cell-epigenome-and-hic-analysis-in-human-mouse-retina). Datasets of human eye 10x multiome and sn-m3C-seq have been uploaded for viewing on the Eye Epigenomics Web Portal (https://epigenome.wustl.edu/EyeEpigenome/).

## Code availability

Code to perform the analyses in this study is accessible at GitHub (https://github.com/yingyuan830/human_retina).

## Supplementary Materials

**The PDF file includes:**

Materials and Methods

Figs. S1 to S10

**Other Supplementary Materials for this manuscript includes:**

Tables S1 to S14

## Materials and Methods

### Nucleus preparation from frozen human retina tissue for Chromium single-cell multiome ATAC and gene expression and snm3C-seq assays

The eye globes were collected from three unrelated human donors (aged 21, 27 and 43 years) with the collaboration of San Diego Eye Bank. The sample processing time starting from eye enucleation to dissection was restricted within 2 hours to identify the macula, a visible yellowish/whiteish indent that is slightly lateral to the optic disk inside the globe. Prior dissection, the eye globe with proper label was collected in a sterile moist chamber. At first the cornea was removed by cutting 2-3 mm around the limbus using a surgical blade. Then the iris and lens were pulled off, with the eye cup remaining, which contained the posterior pole. After removing enough vitreous and aqueous fluid the temporal region can be located by two main blood vessels that curl around the macula. Using a sterile 3mm skin biopsy punch, the macula was collected. Sections were made for the other regions, temporal, nasal, superior and inferior. For all these regions, the retina and RPE tissue were separated and stored immediately in liquid nitrogen.

Retina tissues were pulverized using a dounce tissue chamber and two pestles (“Loose” and “Tight”) on ice for each sample. Pulverized retina tissues were resuspended in 1ml of chilled NIM-DP-L buffer (0.25 M sucrose, 25 mM KCl, 5 mM MgCl2, 10 mM Tris-HCl pH 7.5, 1 mM DTT, 1x protease inhibitor (Pierce), 1U μl^-1^ recombinant RNase inhibitor (Promega, PAN2515) and 0.1% Triton X-100). Tissue was dounce-homogenized with a loose pestle (5-10 strokes) followed by a tight pestle (15-25 strokes) or until the solution was uniform. Nuclei were filtered using a 30 μm CellTrics filter (Sysmex, 04-0042-2316) into a LoBind tube (Eppendorf, 22431021) and flow-through was equally split for 10x mutiome and snm3C-seq assays.

For snm3C-seq, nuclei were pelleted (at 1000 rcf, for 10 min at 4°C; Eppendorf, 5920R) then resuspended in 1mL PBS with 2% formaldehyde for crosslinking for 10 minutes at room temperature on a rotator. 2.5M glycine was added to achieve 0.22M glycine and incubated for 5 minutes at room temperature on a rotator. Nuclei were pelleted 1000 rcf, for 10 min at 4°C then resuspended in 1mL of cold PBS. Nuclei were then pelleted at 2,500 x g for 5 min at 4°C. All supernatant was removed, and pellets were snap frozen in liquid nitrogen and stored at -80°C.

For 10x multiome assays, nuclei were centrifuged 1000 rcf, for 10 min at 4°C then the pellet was resuspended in 1ml NIM-DP buffer (0.25 M sucrose, 25 mM KCl, 5mM MgCl2, 10 mM Tris-HCl pH 7.5, 1mM DTT, 1x protease inhibitor, 1U μl^-1^ recombinant RNase inhibitor) and pelleted (1000 rcf, 10 min at 4°C). Pelleted nuclei were resuspended in 400 μl 2 μM 7-AAD (Invitrogen, A1310) in sort buffer (1mM EDTA, 1U μl^-1^ recombinant RNase inhibitor, 1x protease inhibitor, 1% fatty acid-free BSA in PBS). A total of 150,000 nuclei was sorted (Sony, SH800S) into a LoBind tube containing a collection buffer (5U μl^-1^ recombinant RNase inhibitor, 1x protease inhibitor, 5% fatty acid-free BSA in PBS). Then, 5x permeabilization buffer (50 mM Tris-HCl pH 7.4, 50 mM NaCl, 15 mM MgCl2, 0.05% Tween-20, 0.05% IGEPAL, 0.005% Digitonin, 5% fatty acid-free BSA in PBS, 5mM DTT, 1U μl^-1^ recombinant RNase inhibitor, 5x protease inhibitor) was added for a final concentration of 1x. Nuclei were incubated in ice for 1 min, then centrifuged (500 rcf, 5 min at 4°C). The supernatant was discarded and 650 μl of wash buffer (10 mM Tris-HCl pH7.4, 10 mM NaCl, 3 mM MgCl2, 0.1% Tween-20, 1% fatty acid-free BSA in PBS, 1mM DTT, 1U μl^-1^ recombinant RNase inhibitor, 1x protease inhibitor) was added without disturbing the pellet followed by centrifuging (500 rcf, 5 min at 4°C). The supernatant was removed, and the pellet was resuspended in 7μl of 1x nucleus buffer (nucleus buffer (10x Genomics), 1mM DTT, 1U μl^-1^ recombinant RNase inhibitor). Nuclei (1μl) were diluted in 1x nucleus buffer, stained with Trypan Blue (Invitrogen, T10282) and counted, In total, 16000-20000 nuclei were used for the tagmentation reaction and controller loading and libraries were generated according to the manufacturer’s recommended protocol (https://www.10xgenomics.com/support/single-cell-multiome-atac-plus-gene-expression). 10x multiome ATAC-seq and RNA-sequencing (RNA-seq) libraries were paired-end sequenced on the illumina NovaSeq X Plus systems to a depth of around 50000 reads per cell for each modality.

### Nucleus preparation from fresh RPE cell lines for bulk ATAC-seq

The hTERT-RPE1 (CRL-4000, ATCC) cells were used to introduce a homozygous and heterozygous TMEM216 c.−69G>A variant by Synthego Corporation (*56*). Synthego Corporation generated two cell lines harboring homozygous TMEM216 c.−69G>A variant (D4 and F6) and along with one heterozygous cell line (G1) harboring TMEM216 c.−69G>A variant.

The freshly collected cells were used from wild type hTERT-RPE1, homozygous TMEM216 c.−69G>A clones D4 and F6 and G1, the heterozygous cell line for bulk ATAC-seq. Each cell line was assayed in two replicates, in total there were eight fresh samples applied in the same experiment batch.

For each fresh cell line sample, permeabilized nuclei were obtained by resuspending cells in 250µL Nuclear Permeabilization Buffer [0.2% IGEPAL-CA630 (I8896, Sigma), 1mM DTT (D9779, Sigma), Protease inhibitor (05056489001, Roche), 5% BSA (A7906, Sigma) in PBS (10010-23, Thermo Fisher Scientific)], and incubating for 5 min on a rotator at 4°C. Nuclei were then pelleted by centrifugation for 5 min at 500 xg at 4°C. The pellet was resuspended in 25µL ice-cold Tagmentation Buffer [33 mM Tris-acetate (pH = 7.8) (BP-152, Thermo Fisher Scientific), 66 mM K-acetate (P5708, Sigma), 11 mM Mg-acetate (M2545, Sigma), 16 % DMF (DX1730, EMD Millipore) in Molecular biology water (46000-CM, Corning)]. An aliquot was then taken and counted by hemocytometer to determine nuclei concentration. Approximately 50,000 nuclei were resuspended in 20µL ice-cold Tagmentation Buffer, and incubated with 1µL Tagmentation enzyme (FC-121-1030; Illumina) at 37 °C for 60 min with shaking 500 rpm. The tagmentated DNA was purified using MinElute PCR purification kit (28004, Qiagen). The libraries were amplified using NEBNext High-Fidelity 2X PCR Master Mix (M0541, NEB) with primer extension at 72°C for 5 min, denaturation at 98°C for 30s, followed by 8 cycles of denaturation at 98°C for 10 s, annealing at 63°C for 30 s and extension at 72°C for 60 s. Amplified libraries were then purified using MinElute PCR purification kit (28004, Qiagen), and two size selection steps were performed using SPRIselect bead (B23317, Beckman Coulter) at 0.55X and 1.5X bead-to-sample volume rations, respectively. The ATAC-seq libraries were paired-end sequenced on the Illumina NextSeq 2000 system to a depth of around 50 M reads per sample.

### Genome assemblies and annotations

Homo sapiens (human) assembly: hg38, GRCh38 annotation: hg38 Gencode v33.

### 10x multiome data processing and clustering

Raw sequencing data were processed using cellranger-arc (10x Genomics), generating single-nucleus RNA-seq (snRNA-seq) UMI count matrices for intronic and exonic reads mapping in the sense direction of a gene.

### Nucleus isolation and FANS

For all snm3C-seq samples, in situ 3C treatment was performed during the nucleus preparation, enabling the capture of the chromatin conformation modality as described previously (*29*). These steps were performed using the Arima-3C BETA Kit (Arima Genomics). The nuclei were isolated and sorted into 384-well plates using previous described methods (*38*). Briefly, nuclei were stained with DRAQ7 (ThermoFisher Scientific, D15106) and then processed for fluorescence-activated nucleus sorting (FANS) using the Sony SH800 cell sorter with single-cell (1 drop single) mode.

### Library preparation and Illumina sequencing

The snm3C-seq samples were prepared according to a previously described library preparation protocol (*29*, *38*). This protocol has been automated using the Beckman Biomek i7 instrument to facilitate large-scale applications. The snm3C-seq libraries were shallow sequenced on the Illumina NovaSeq 2000 to 2-10M reads to check quality. Successful libraries were deeply sequenced on the NovaSeq X Plus instrument, using one lane of 25B flowcell per 4 384-well plates and using 150 bp paired-end mode.

### Nucleus preparation from frozen mouse retina tissue

The flash frozen retinal tissue along with RPE and choroid was harvested from 2.5month, 5month old mice. Each age point contained two biological replicates. Nuclei preparation was adapted from Lacar et al. 2016 (PMID: 27090946). Flash frozen mouse retina was resuspended in 1 mL of douncing buffer consisting of 0.25M sucrose (S1888, Sigma), 25mM KCl (AM9610G, Invitrogen), 5mM MgCl2 (194698, Mp Biomedicals Inc.), 10 mM Tris-HCl pH 7.5 (15567027, Thermo Fisher Scientific), 1mM DTT (D9779, Sigma), 1X protease inhibitor (05056489001, Roche), 0.1% Triton X-100 (T8787-100ML, Sigma), and 0.5 U/ul RNasin RNase inhibitor (PAN21110, Promega) in molecular biology grade water (46000-CM, Corning). Frozen tissue was then transferred to a dounce homogenizer on ice and dounced 25 times with a loose plunger and 25 times with a tight plunger. Suspension was then passed through a Celltrix 30 μM filter (04-004-2326, Sysmex) and washed with 300 mL of douncing buffer before being centrifuged for 10 minutes at 1000 rcf in a swinging bucket centrifuge at 4°C with run setting 3/3. Pellet was then washed with an additional 1 mL of douncing buffer without Triton X-100, and split for single-nucleus ATAC-seq and single-nucleus RNA-seq in a ratio of 700 mL /300 mL.

### Single-nucleus ATAC-seq assay of mouse retina

Combinatorial barcoding single-nucleus ATAC-seq assay was performed as previously described (*63–65*). Nuclei from dissociated tissue were permeabilized in 1 mL of permeabilization buffer (5% BSA, 0.2% IGEPAL-CA630, 1 mM DTT, 1x cOmplete EDTA-free protease inhibitor in PBS), pipette mixed, and incubated on ice for 10 minutes. Nuclei were then centrifuged at 500 rcf for 5 minutes and the supernatant was removed. Nuclei were resuspended in 500 µL high salt tagmentation buffer (36.3 mM Tris-acetate (pH = 7.8), 72.6 mM potassium-acetate, 11 mM Mg-acetate, 17.6% DMF) and counted using a hemocytometer. Concentration was adjusted to 2000 nuclei/9 µL, and 2000 nuclei were dispensed into each well of one 96-well plate. For tagmentation, 1 µL barcoded Tn5 transposomes (*66*) was added using a BenchSmart 96 (Mettler Toledo), mixed five times and incubated for 60 min at 37°C with shaking (500 rpm). To inhibit the Tn5 reaction, 10 µL of 40 mM EDTA were added to each well with a BenchSmart 96 (Mettler Toledo) and the plate was incubated at 37°C for 15 min with shaking (500 rpm). Next, 10 µL 3x sort buffer (2% BSA, 2 mM EDTA in PBS) was added using a BenchSmart 96 (Mettler Toledo). All wells were combined into a FACS tube and stained with 3 µM Draq7 (Cell Signaling). Using a SH800 (Sony), 20 2n nuclei were sorted per well into eight 96-well plates (total of 768 wells) containing 10.5 µL EB (25 pmol) primer i7, 25 pmol primer i5, 200 ng BSA (Sigma). Preparation of sort plates and all downstream pipetting steps were performed on a Biomek i7 Automated Workstation (Beckman Coulter). After addition of 1 µL 0.2% SDS, samples were incubated at 55°C for 7 min with shaking (500 rpm). 1 µL 12.5% Triton-X was added to each well to quench the SDS. Next, 12.5 µL NEBNext High-Fidelity 2 × PCR Master Mix (NEB) were added and samples were PCR-amplified (72°C 5 min, 98°C 30 s, (98°C 10 s, 63°C 30 s, 72°C 60 s)×12 cycles, held at 12°C). After PCR, all wells were combined. Libraries were purified according to the MinElute PCR Purification Kit manual (Qiagen) using a vacuum manifold (QIAvac 24 plus, Qiagen) and size selection was performed with SPRI Beads (Beckmann Coulter, 0.55x and 1.5x). Libraries were purified one more time with SPRI Beads (Beckmann Coulter, 1.5x). Libraries were quantified using a Qubit fluorometer (Life technologies) and the nucleosomal pattern was verified using a Tapestation (High Sensitivity D1000, Agilent). The library was sequenced on a NovaSeq6000 or NextSeq500 sequencer (Illumina) using custom sequencing primers with following read lengths: 50 + 10 + 12 + 50 (Read1 + Index1 + Index2 + Read2).

### Single-nucleus RNA-seq assay of mouse retina

Single nucleus RNA-seq was carried out using the Droplet-based Chromium Single-Cell 3’ solution (10x Genomics, v3 chemistry) (*67*). Nuclei from dissociated tissue were centrifuged at 1000 rcf for 10 minutes. Supernatant was then discarded, and the pellet was resuspended in 500 mL of sort buffer consisting of 1 mM EDTA (15575020, Invitrogen), 0.5 U/ul RNasin, and 1% BSA in PBS and split for single-nucleus RNA-seq (snRNA-seq) and single-nucleus ATAC-seq (snATAC-seq). Nuclei from dissociated tissue were stained with 3 μM DRAQ7 (#7406S, Cell Signaling Technology). Nuclei were then incubated on ice for 10 minutes and ∼50,000-75,000 nuclei were sorted using a 100 μm chip in an SH800 sorter (Sony) into 50 ul of collection buffer consisting of 2.5 U/ul RNasin and 5% BSA in PBS. Samples were then centrifuged for 15 minutes at 1000 rcf; the supernatant was removed, leaving behind ∼20 mL, and an additional 25 ul of reaction buffer consisting of 0.5 U/ul RNasin and 1% BSA in PBS was added for a total volume of ∼45-50 ul. Nuclei were visually inspected and manually counted using a hemocytometer before loading 15,000 onto a Chromium Controller for 10x GEM generation in the Single Cell 3’ v3.1 kit (1000268, 10x Genomics). Libraries were generated using the Chromium Single-Cell 3′ Library Construction Kit v3 (10x Genomics, 1000075) with the Chromium Single-Cell B Chip Kit (10x Genomics, 1000153) and the Chromium i7 Multiplex Kit for sample indexing (10x Genomics, 120262) according to manufacturer specifications. CDNA was amplified for 12 PCR cycles. SPRISelect reagent (Beckman Coulter, B23319) was used for size selection and clean-up steps.

Final library concentration was assessed by Qubit dsDNA HS Assay Kit (Thermo-Fisher Scientific) and fragment size was checked using Tapestation High Sensitivity D1000 (Agilent) to ensure that fragment sizes were normally distributed about 500 bp. Libraries were sequenced using the NextSeq550 and a NovaSeq6000 (Illumina) with these read lengths: 28 + 10 + 10 + 91 (Read1 + Index1 + Index2 + Read2).

## Data preprocessing

### Identification of open chromatin regions from snATAC-seq data of each cell cluster

Peak calling was performed on the Tn5-corrected single-base insertions using the MACS2 with these parameters on pseudo bulk ATAC-seq fragments: –shift -75 –extsize 150 –nomodel – call-summits –SPMR -q 0.05 (for the cluster’s cell number is larger than 1000) or 0.1 (for the cluster’s cell number is less than 1000). We extended peak summits by 250 bp on either side to a final width of 501 bp for merging and downstream analysis. Since the number of peaks called in each cell type is related to the sequence depth, which is highly variable due to differences in cell type abundance, we converted MACS2 peak scores (-log10[q]) to scores per million. Peaks with a score per million of >=2 were retained for each cell type. We also filtered human and mouse peaks by removing those with ECODE blacklist regions of hg38 or mm10.

### Identification of top marker genes of snRNA-seq clusters

We used the FindAllMarkers function of Seurat (*31*) to identify the positive and negative markers of an annotated single cluster, compared to all other cell types. In brief, we performed a logistic regression-based method to identify differentially expressed (DE) genes. The reported top 50 markers displayed in the heat map are ranked based on their average log2 fold change (avg_log2FC), with metrics (logfc.threshold = 0.25, min.pct = 0.25) used to determine the most significant markers.

### Identification of chromatin accessibility level at cCREs

For each cell type, we normalized peak accessibility in each cluster to log2[CPM+1] quantified for ATAC-seq peaks.

### Identification of gene expression level

For each cell type, we normalized gene expression in each cluster to log2[CPM+1] quantified for human genes (the counts are UMI here).

### Identification of significant cell type specific cCREs or genes

Based on the known identified cCREs chromatin accessibility level and gene expression level matrix (rows as cCREs or genes, columns as cell types), we further find the significant cell type specific cCREs or genes. First, we calculated the z score of each row with (row_data - mean of row_data)) / sd of row_data; then identify the max z-score of each row and compute the p-value for max z-score using the cumulative distribution function (CDF) of the standard normal distribution: 2 * (1 - pnorm(abs(max_z_score))) as the p-value for each row, this provides a two-tailed test; finally we filtered and selected the rows with p_value <= 0.05 to be classified as significant cell type specific cCREs or genes.

### Mapping of snm3C-seq

Reads from FASTQ files were mapped using YAP (Yet Another Pipeline) software (cemba-data v1.6.9), as previously described (*44*). First, FASTQ files were demultiplexed for each cell barcode. Next, reads were assessed for quality, then two-pass mapping was performed with bismark (v0.20, with bowtie2 v2.3). BAM file processing and QC was performed using samtools (v1.9) and Picard (v3.0.0). Chromatin contacts were called and methylome profiles were generated using Allcools (v1.0.23). All reads were mapped to the hg38 genome assembly.

### Quality control of snm3C-seq

We filtered for high quality cells based on DNA methylation by requiring mapping rate for both reads R1 and R2 > 0.5; total final reads > 100,000; overall mCCC level < 0.03; overall mCH level < 0.2; and overall mCG level > 0.5.

### Methylome clustering analysis

Single-cell DNA methylome profiles were stored in the ‘all cytosine’ (ALLC) format, a tab-separated table compressed and indexed by bgzip/tabix. We used the generate-dataset command in the ALLCools package to generate a methylome cell-by-feature tensor dataset (MCDS) in Zarr format. We applied the integration with the companion 10x multiome RNA-seq dataset:

(1) Basic feature filtering: exclude regions in chrX, chrY, chrM and chrL, and ENCODE blacklist.
(2) Apply principal component analysis (PCA) on merged data of mC and RNA
(3) Integrate the data by method similar to Seurat (*68*)
(4) Transfer the annotation of labeled RNA-seq to mC-seq data which belongs to the same cluster with RNA-seq data to label the cells of mC clusters.

### Cluster-level DNA methylome analysis

After the integration with RNA-seq profile, we merged the single-cell ALLC files into pseudo-bulk level using the ALLCools merge-allc command. Next we performed DMR calling using methylpy as described previously (*40*). Briefly, we first calculated CpG differentially methylated sites using a permutation-based root mean square test. The base calls of each pair of CpG sites were added before analysis. We then merged the differentially methylated sites into DMR if they are within 500 bp.

### Cell- and cluster-level 3D genome analysis

#### Generating the chromatin contact matrix and imputation

After snm3C-seq mapping, we used the cis long-range contacts (contact anchors distance > 2500 bp) and trans contacts to generate single-cell raw chromatin contact matrices at three genome resolutions: chromosome 100 kb resolution for the chromatin compartment analysis; 25 kb bin resolution for the chromatin domain boundary analysis; 20 kb resolution for the chromatin loop or dot analysis. The raw cell-level contact matrices are stored in HDF5-based scool format. We then used the scHiCluster package (v.1.3.2) (*69*) to perform contact matrix imputation. The scHiCluster imputes the sparse single-cell matrix in two steps: (1) Gaussian convolution (pad = 1) (2) apply a random walk with restart algorithm on the convoluted matrix. The imputation was performed on each cis matrix (intrachromosomal matrix) of each cell. For 100 kb matrices, the whole chromosome was imputed; for 25 kb matrices, we imputed contacts within 10.05 Mb; for 10 kb matrices, we imputed contacts within 5.05 Mb. The imputed matrices for each cell were stored in cool format. For most of the following analyses, cell matrices were aggregated into cell groups identified in the previous section. These pseudo-bulk matrices are concatenated into a tensor called CoolDS, and stored in the Zarr format.

### Domain boundary analysis

We applied the imputed cell-level contact matrics at 25 kb resolution to identify the domain boundaries within each cell using the TopDom algorithm (*70*). We also filtered out the boundaries that overlap with ENCODE blacklist v2. We used cooltools (v.0.5.1) to call cluster-level boundaries and domains with 25 kb resolution matrices. A sliding window of 500 kb was used to compute the insulation score of each bin; the bins with boundary strength > 0.1 were selected as domain boundaries.

### Identification of orthologous sequence elements across species

We identified orthologous sequences for the human cis-regulatory region in mice using liftOver (*71*). For each human ATAC peak, we performed liftOver to the mouse genome with a requirement of 50% retained sequence identity (minMatch = 0.5). Any region that couldn’t be lifted to any of the other profiled species was identified as human specific. For ATAC peaks (500 bp), we retained only orthologous elements that are 1 kb or less to the lifted-over genome. We next performed liftOver from the identified orthologous sequence back to the human sequence. We retained all sequences that mapped back to the same peak identity as ‘human conserved’ between human and mouse.

### Identification of DMRs

We performed DMR calling using the call_dms and call_dmr functions of ALLCools (*38*). In brief, we first calculated CpG differential methylated sites using a permutation-based root mean square test (*72*), the base calls of each pair of CpG sites were combined before analysis. We then merged the differential methylated site into a DMR if they were within 250 bp and had PCC > 0.3 across samples. We applied the DMR calling framework across each cell type within retina and macula, RPE tissue.

### Paired cell type specificity of ATAC peaks, genes and DMRs

For each ATAC peak, gene, and DMR, we set the ATAC peak as the cell type specific peak if the log2[CPM+1] is highest for one cell type, then we showed the paired target gene from the ABC model prediction. For DMRs, we transformed quantifications to 1-the methylation level in each cell type, the corresponding DMRs are also plotted to check the consistency.

### GO enrichment analysis

We performed GO enrichment analysis using the Enrichr module(*73*) in GSEApy(*74*). For each gene set, we used the GO biological process 2023. We performed using the different appropriate background set according to the different cases: the background set for across species gene differences analysis was all genes expressed in human and mouse for a specific cell type; for ABC target genes, the background set was all human genes called as having an ABC enhancer; for cell type marker genes, the background set was all marker genes from all the cell types.

### Identification of species-biased gene activity

Based on the list of one-to-one orthologous genes across human and mouse, we performed differential expression analysis on pseudo bulk count profiles for each cell type using edgeR (v.3.36.0) (*75*). To account for multiple comparisons, we nominated an FDR of 0.001 as a threshold, besides, we required our differentially expressed genes to meet a minimum fold change of 2. After applying these criteria, we further identified biased genes for each species, for each cell type in each species, we identified biased genes as a gene that was significantly upregulated in that cell type compared with in each other species.

### TF motif scanning

We performed both de novo and known motif enrichment analysis using Homer (v4.11.1) (*76*). For cCREs in the consensus list, we scanned a region of ± 250 bp around the summit of the element. Randomly selected background regions are used for motif discovery. We adopted motifs with q value <= 0.05 finally.

### Identifying putative enhancer-gene pairs with the ABC model

We used the ABC model (*46*) to identify putative enhancer genelinks in each species. Briefly, the ABC model uses normalized contact frequencies from HiC data, along with a measure of enhancer activity, to predict putative enhancer-gene pairs. For each cell type, we ran the ABC model using the default parameters, providing normalized HiC matrixes at 10 kb resolution, ATAC chromatin accessibility BAM files and a list of ATAC peaks identified in that same cell type. Predictions with an ABC score greater or equal to 0.02 were considered positive and used for the downstream analysis.

### Mouse sequence data analysis

For the mouse snRNA-seq data analysis, we first used CellBender (*78*) to remove ambient and background RNA, then applied Seurat to do QC filtering, clustering, annotation and other processing. The QC metrics were applied on nFeature_RNA (100–7000) and percent.mt, besides, DoubletFinder was used to remove classified doublets. Next, the datasets were merged and normalized through SCTransform, we also performed batch correction via Harmony which was wrapped in Seurat. During the clustering, we set dimensions as 10 and resolution as 0.5. Finally, the clusters were annotated by using the cell type specific marker genes searched from published literature, we identified 13 cell types totalling 53299 nuclei from this modality.

As for the snATAC-seq data analysis, we first used Sinto (https://github.com/timoast/sinto) to create the sciATAC-seq fragment files, then applied Signac to perform QC filtering (the metrics are TSS.enrichment > 5, Unique_usable_reads > 1000; besides, Scrublet was used to calculate, predict and remove the doublets) and other processing. After merging different samples’ Seurat object to be one, the merged object was normalized through TF-IDF normalization, and we also performed batch correction via Harmony (*79*). During the clustering, we set dimensions as 2:35 and resolution as 1.2. Finally, the clusters were annotated by label transferring from snRNA-seq onto sciATAC-seq clusters, we identified 13 cell types totalling 51220 nuclei from this modality.

### Deep learning model for sequence prediction

We adopted the Basenji (*55*) neural network architecture to predict open chromatin accessibility. We used the same model structure with minor modifications to do the raw prediction, replacement mutation prediction and deletion mutation prediction.

We separated the datasets to be training (80%) and validation (20%) data randomly. The models were trained with a workstation with two NVIDIA GeForce RTX 4090 cards (each one with 24GB GPU memory). We save the model with the highest epoch validation pearsonR during the training and use it for the prediction of mutation.

### GWAS variant enrichment analysis

We obtained GWAS summary statistics for quantitative traits. We prepared summary statistics to the standard format for linkage disequilibrium score regression. We used all union cCREs from each cell type of the retina and macula tissue as the background control set for the retina cell type computation, similar for the RPE tissue cell types. For each trait, we used cell-type-specific linkage disequilibrium score regression (https://github.com/bulik/ldsc) to estimate the enrichment coefficient of each annotation jointly with the background control (*77*).

**Fig. S1.**
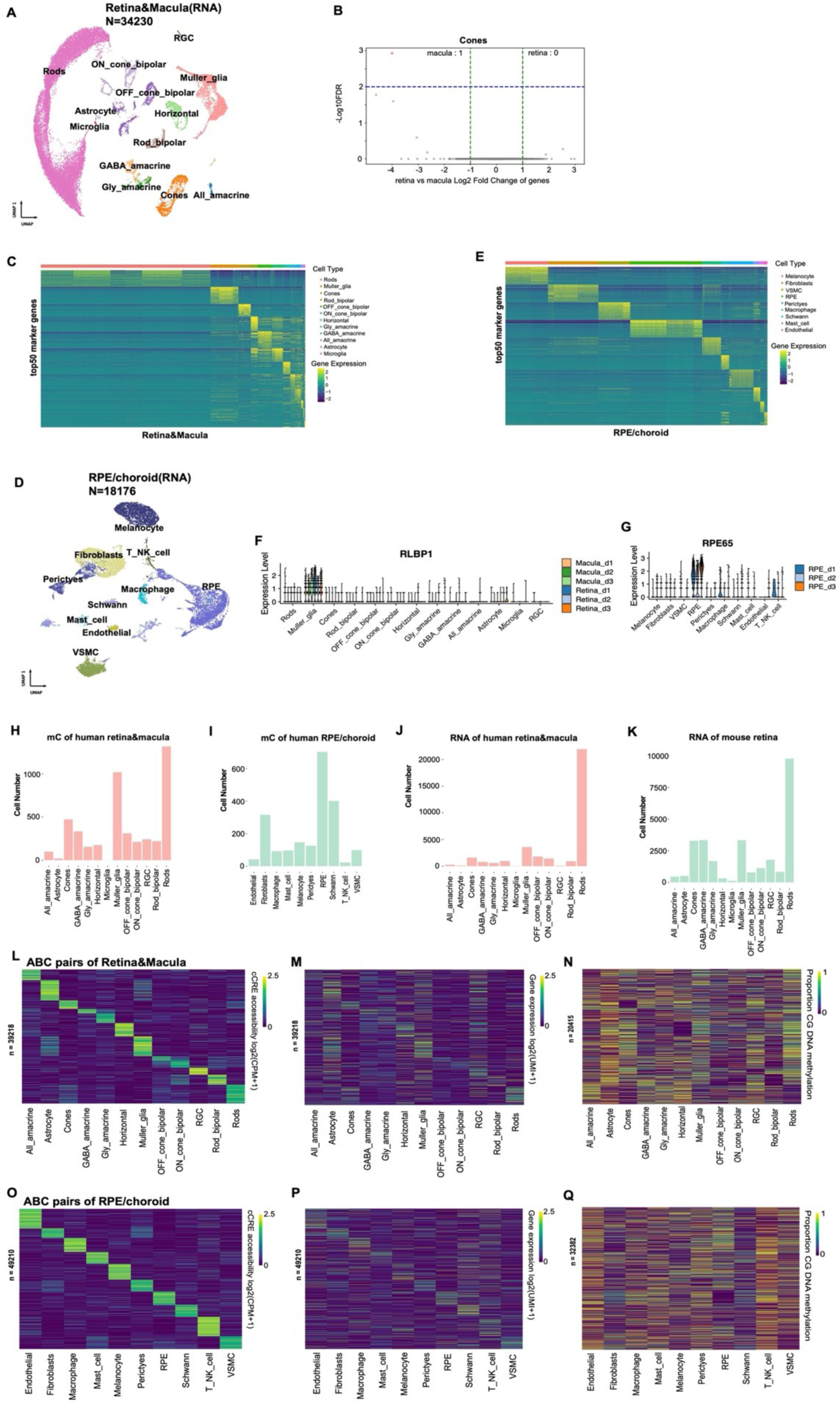
Single-cell multimodal analysis of human retina, macula, RPE/choroid tissues. (**A**) UMAP embedding of 10x multiome RNA-seq data with annotation for human retina and macula cell types. (**B**) Differential gene expression between Cones of human retina and macula. (**C**) Heatmap of top 50 marker gene expression levels for each cell type in retina and macula tissues. **(D)** UMAP embedding of 10x multiome RNA-seq with cell type annotation from human RPE/choroid tissue. (**E**) Heatmap of top 50 marker gene expression levels for each cell type in RPE/choroid tissue. (**F**) Marker gene of Muller glia (*RLBP1*) expression level across different donors for each cell type in retina and macula tissues. **(G)** Marker gene of RPE (*RPE65*) expression level across different donors for each cell type in RPE/choroid tissue. (**H**) Bar plot of cell numbers of each cell type of mC for human retina and macula tissues. (**I**) Bar plot of cell numbers of each cell type of mC for human RPE/choroid tissue. (**J**) Bar plot of cell numbers of each cell type of RNA for human retina and macula tissues. (**K**) Bar plot of cell numbers of each cell type of RNA for mouse retina. (**L**) Heatmap showing chromatin accessibility of cCREs for each cell type of human retina and macula tissues (the peak of peak-gene pairs predicted by ABC model). CPM, counts per million. Each CRE is ordered by the cell type with the highest accessibility level. (**M**) Heatmap showing the expression of paired genes for each cell type of human retina and macula tissues (the gene of peak-gene pairs predicted by ABC model). UMI, unique molecular identifier, here the UMI is processed with the CPM method. (**N**) Heatmap showing the DMRs corresponding to the peak of peak-gene pairs predicted for each cell type of human retina and macula tissues. (**O**) Heatmap showing chromatin accessibility of cCREs for each cell type of human RPE/choroid tissue (the peak of peak-gene pairs predicted by ABC model). (**P**) Heatmap showing the expression of paired genes for each cell type of human RPE/choroid tissue (the gene of peak-gene pairs predicted by ABC model). (**Q**) Heatmap showing the DMRs for each cell type of human RPE/choroid tissue (the DMR is corresponding to the peak of peak-gene pairs predicted by the ABC model).

**Fig. S2.**
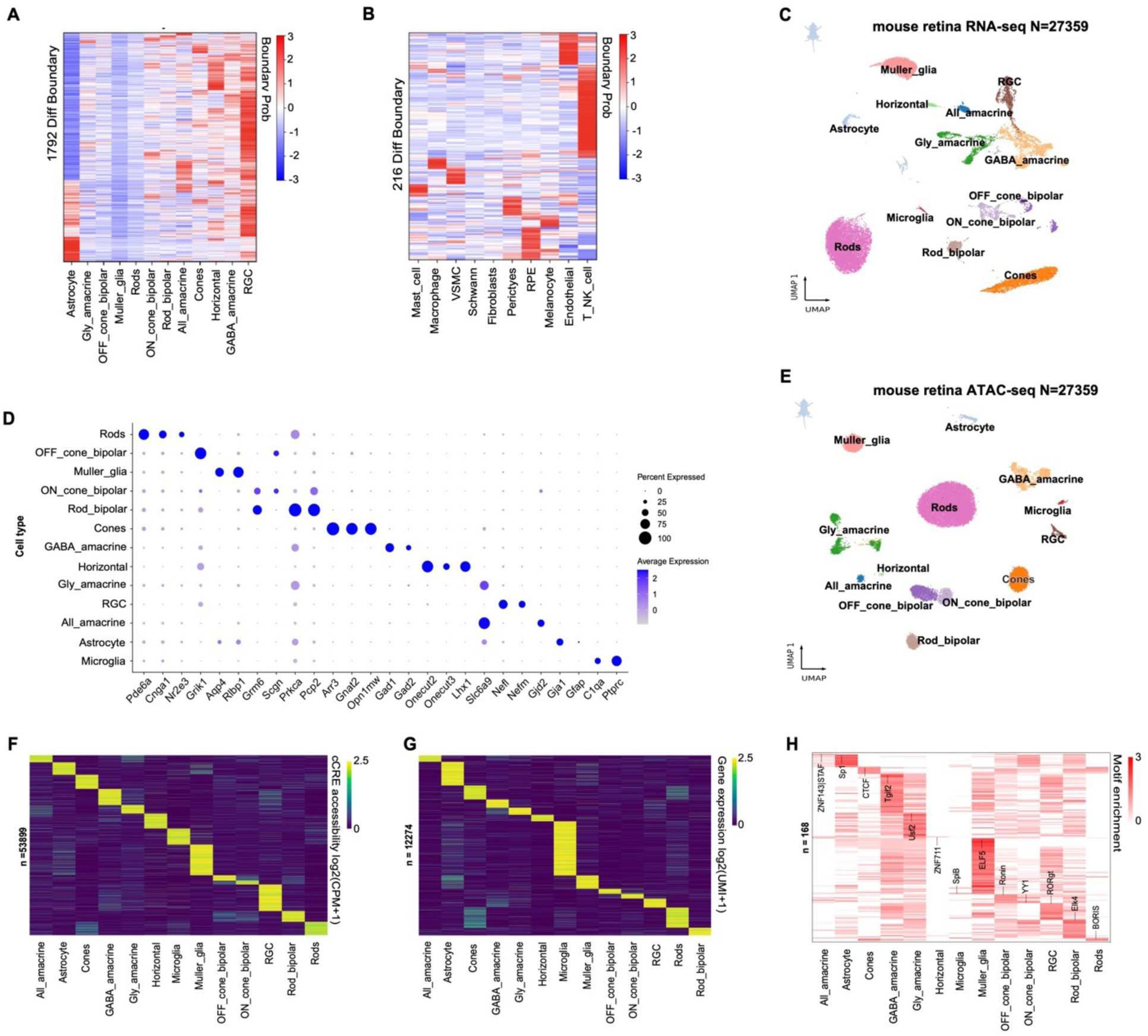
Chromatin domains in human retina cell types and single cell analysis of mouse retina. (**A**) Heatmaps of boundary probability of 1792 identified different TAD boundaries in each cell type of retina and macula tissues. The color bar shows the row-wise z-score normalization value of the boundary probability, red represents higher probability, blue represents lower probability. (**B**) Heatmaps of boundary probability of 216 identified different TAD boundaries in each cell type of RPE tissue. (**C**) UMAP embedding of 10x snRNA-seq data with cell type annotation for mouse retina. **(D)** Dot plot visualizing the normalized RNA expression of selected marker genes by cell type of mouse retina tissue. **(E)** UMAP embedding of 10x snATAC-seq data with cell type annotation from mouse retina. (**F**) Heatmap showing chromatin accessibility of significant cell-type specific cCREs of mouse retina. CPM, counts per million. Each CRE is ordered by the cell type with the highest accessibility level. (**G**) Heatmap showing expression of significant cell-type specific gene of mouse retina. UMI, unique molecular identifier, here the UMI is processed with the CPM method. (**H**) Enrichment of HOMER known TF motifs in cCRE regions of mouse retina.

**Fig. S3.**
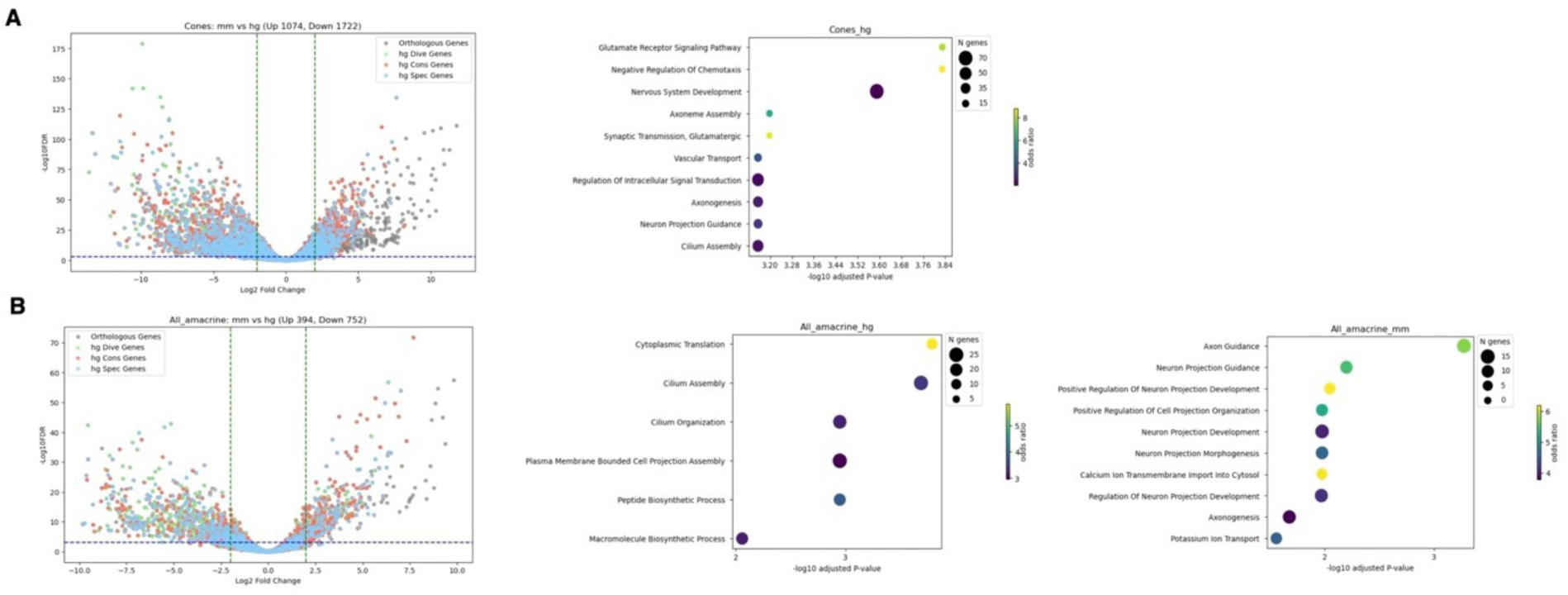
Differential gene expression between human and mouse retinal cell types. (**A-B**) Volcano plot showing the differential gene expression between human and mouse Cones cells (A) or All amacrine cells (B), along with most significant GO terms associated with differentially expressed genes. Gray dots represent all the orthologous genes across human and mouse in this cell type; green dots represent the genes paired with human divergent cCREs; red dots represent the genes paired with human conserved cCREs; blue dots represent the genes paired with human specific cCREs.

**Fig. S4.**
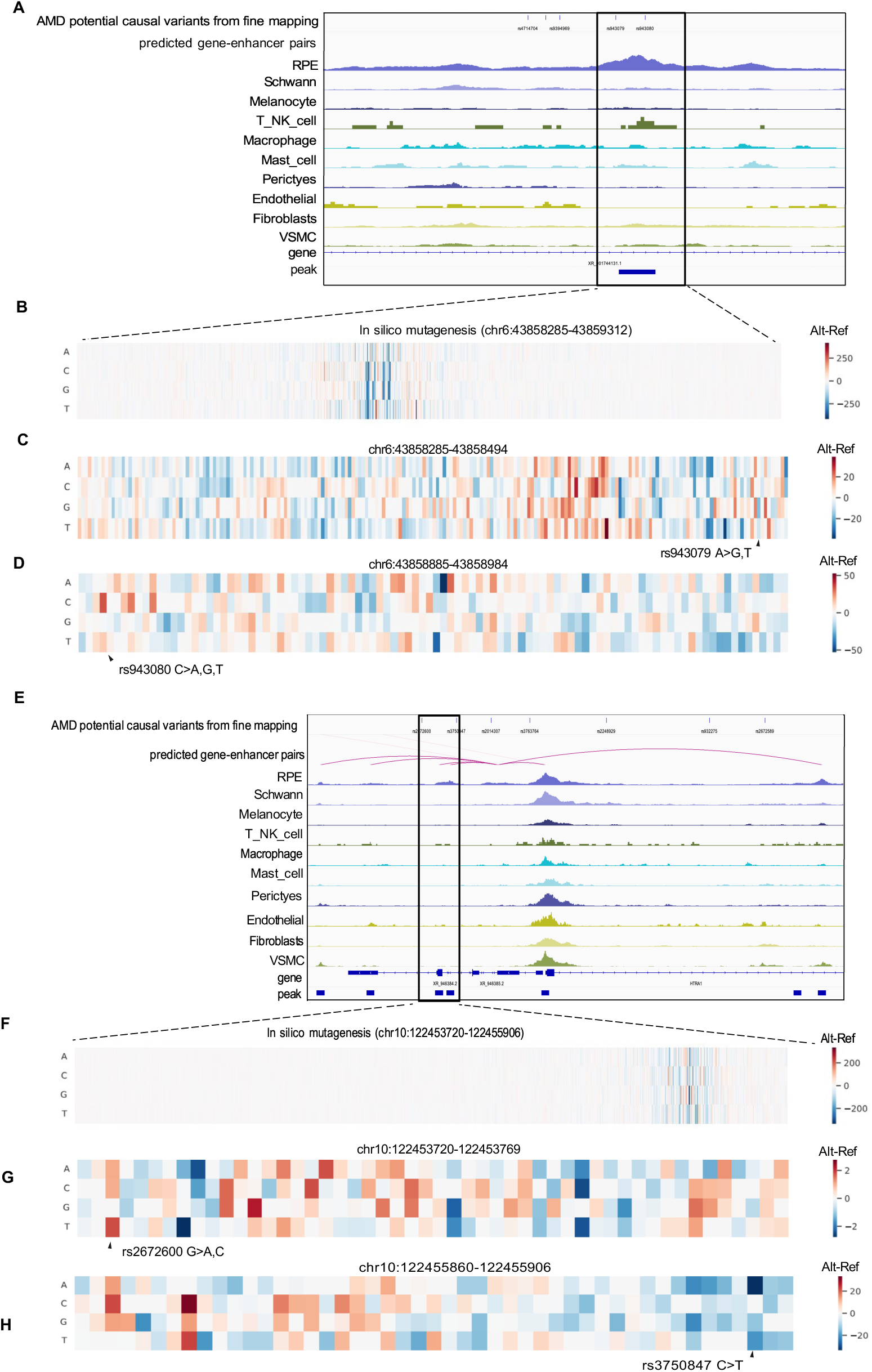
The Deep learning model helps to interpret mutation effects on noncoding risk variants of ocular disorder and traits for the same cell type. (**A and E**) Fine mapping of AMD trait and causal risk variants in different categories of cCREs from cell types of RPE tissue. Genome browser tracks (GRCh38) display chromatin accessibility profiles from snATAC-seq, magenta arcs represent the predicted pairs of enhancer and gene from ABC model in RPE cell type. (**B**) In silico nucleotide mutagenesis influenced the prediction of accessibility within region chr6: 43858285-43859312 (0-based) of RPE cell type. Larger signals (deeper red) represent a higher accessibility prediction on altered sequence, lower signals (deeper blue) represent lower accessibility on altered sequence. (**C**) Zoom in in silico nucleotide mutagenesis within region chr6: 43858285-43858494 of RPE cell type. Lower accessibility predicted on the enhancer with risk variant rs943079 A>G,T. (**D**) Zoom in in silico nucleotide mutagenesis within region chr6: 43858885-43858984 of RPE cell type. Higher accessibility predicted on the enhancer with risk variant rs943080 C>A. (**F**) In silico nucleotide mutagenesis influenced the prediction of accessibility within region chr10: 122453720-122455906 (0-based) of RPE cell type. (**G**) Zoom in in silico nucleotide mutagenesis within region chr10: 122453720-122453769 of RPE cell type. Higher accessibility predicted on the enhancer near to risk variant rs2672600 G>A, C. (**H**) Zoom in in silico nucleotide mutagenesis within region chr10: 122455860-122455906 of RPE cell type. Lower accessibility predicted on the enhancer near to risk variant rs3750847 C>T.

**Fig. S5.**
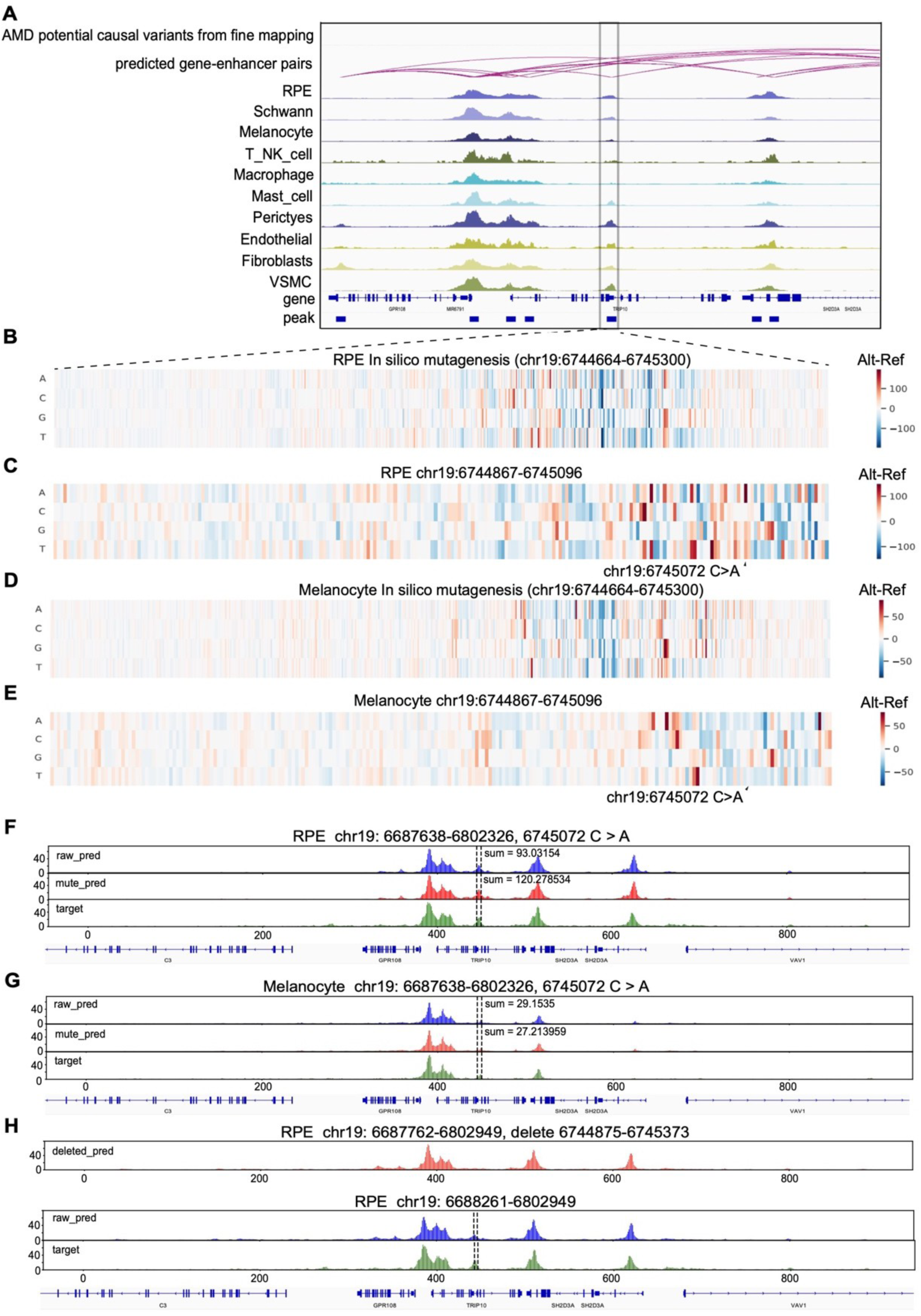
The Deep learning model helps to interpret mutation effects on noncoding risk variants of ocular disorder and traits for different cell types. (**A**) Fine mapping of AMD trait and causal risk variants in different categories of cCREs from cell types of RPE tissue. Genome browser tracks (GRCh38) display chromatin accessibility profiles from snATAC-seq, magenta arcs represent the predicted pairs of enhancer and gene from ABC model in RPE cell type. (**B** and **D**) In silico nucleotide mutagenesis influenced the prediction of accessibility within region chr19: 6744664-6745300 (0-based) of RPE cell type (B) and Melanocyte (D). Larger signals (deeper red) represent a higher accessibility prediction on altered sequence, lower signals (deeper blue) represent lower accessibility on altered sequence. (**C** and **E**) Zoom in in silico nucleotide mutagenesis within region chr19: 6744867-6745096 of RPE cell type (C) and Melanocyte (E). Higher accessibility predicted on the *TRIP10* enhancer with potential variant loci chr19 6745072 C>A for RPE and lower accessibility for Melanocyte. (**F** and **G**) Chromatin accessibility at *TRIP10* enhancer loci predicted in human RPE cell type (F) and Melanocyte (G). Green represents the raw target track, blue represents raw predicted track, red represents predicted track after mutation. (**H**) Chromatin accessibility after deleting TRIP10 enhancer loci predicted in human RPE cell type. Red represents predicted track after deletion mutation.

**Fig. S6.**
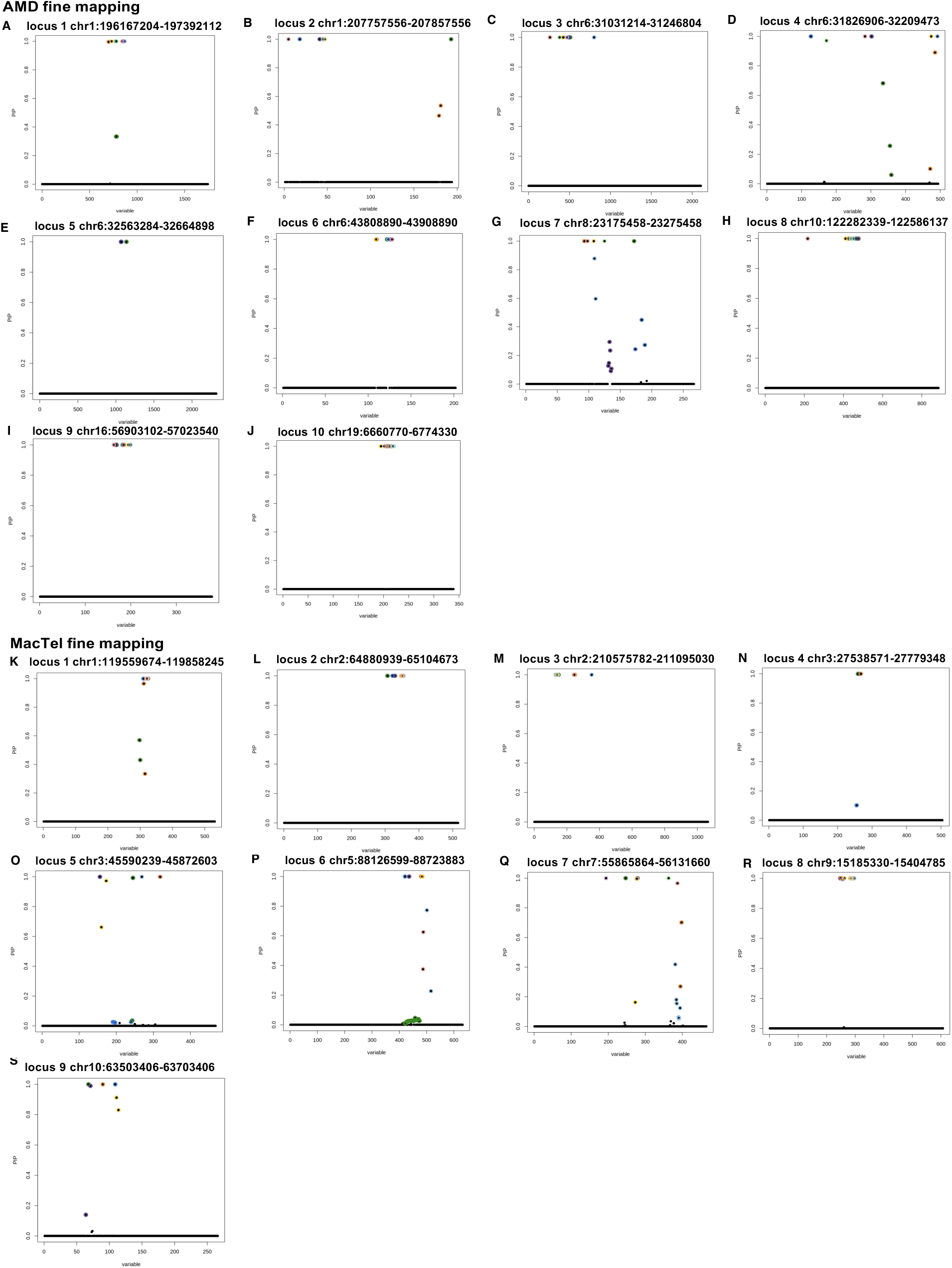
Loci of AMD trait and MacTel trait from fine mapping. (**A-J**) 10 found loci of AMD trait from fine mapping. PIP: Posterior Inclusion Probability. Colored spots represent causal SNPs in the region. (**K-S**) 9 found loci of MacTel trait from fine mapping.

**Fig. S7.**
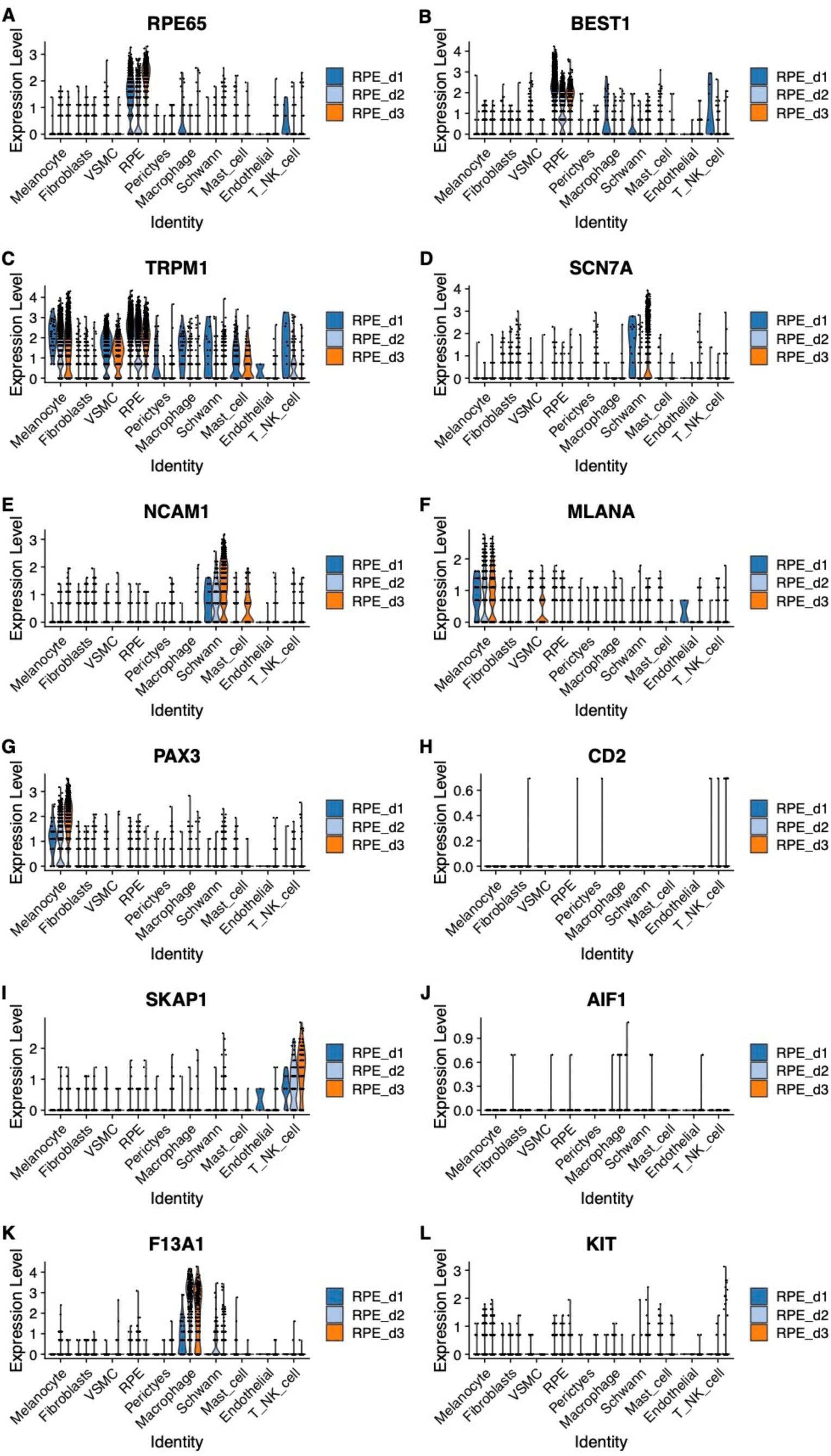
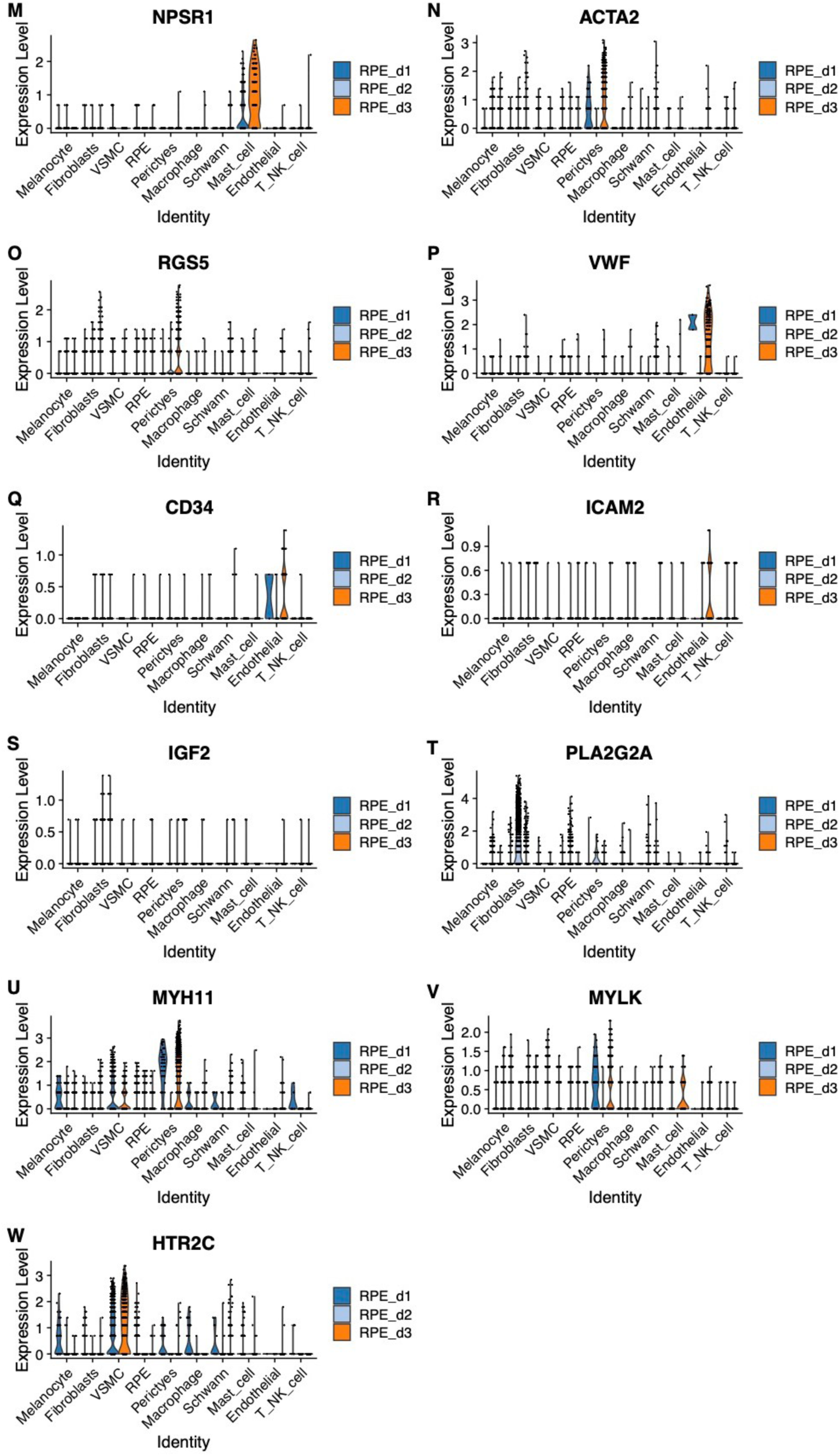
Marker gene expression level across different donors for each cell type in RPE/choroid tissue. **(A)** Marker gene RPE65 **(B)** BEST1 **(C)** TRPM1 **(D)** SCN7A **(E)** NCAM1 **(F)** MLANA **(G)** PAX3 **(H)** CD2 **(I)** SKAP1 **(J)** AIF1 **(K)** F13A1 **(L)** KIT **(M)** NPSR1 **(N)** ACTA2 **(O)** RGS5 **(P)** VWF **(Q)** CD34 **(R)** ICAM2 **(S)** IGF2 **(T)** PLA2G2A **(U)** MYH11 **(V)** MYLK **(W)** HTR2C. The values of gene expression were normalized with function SCTransform() of Seurat, the expression level shows the normalized results of gene expression for better visualization and comparison.

**Fig. S8.**
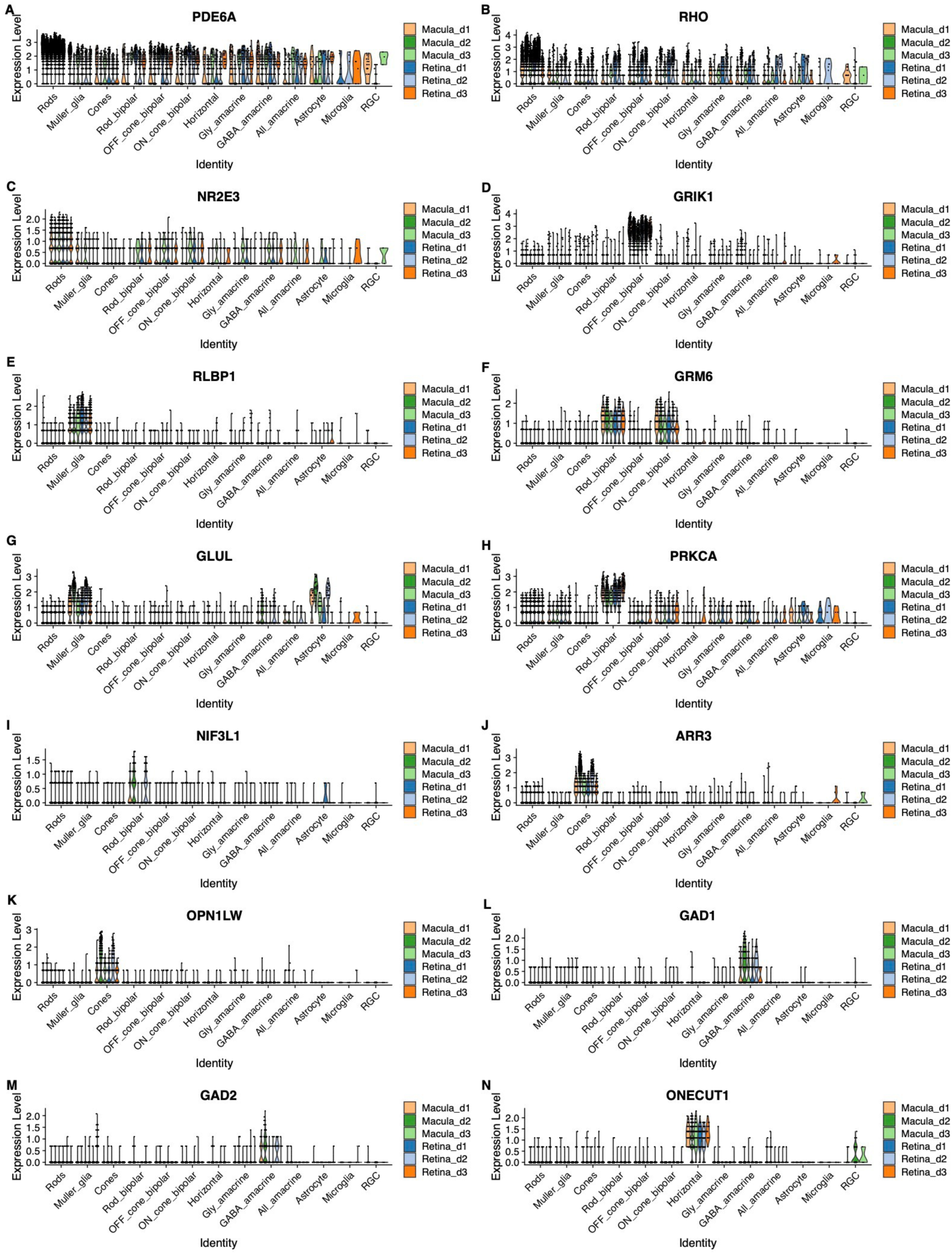
Marker gene expression level across different donors for each cell type in retina and macula tissue. **(A)** Marker gene PDE6A **(B)** RHO **(C)** NR2E3 **(D)** GRIK1 **(E)** RLBP1 **(F)** GRM6 **(G)** GLUL **(H)** PRKCA **(I)** NIF3L1 **(J)** ARR3 **(K)** OPN1LW **(L)** GAD1 **(M)** GAD2 **(N)** ONECUT1 **(O)** ONECUT2 **(P)** LHX1 **(Q)** SLC6A9 **(R)** NEFL **(S)** NEFM **(T)** SLC17A6 **(U)** GJD2 **(V)** CALB2 **(W)** GFAP **(X)** PAX2 **(Y)** HLA-DRA **(Z)** C1QA. The values of gene expression were normalized with function SCTransform() of Seurat, the expression level shows the normalized results of gene expression for better visualization and comparison.

**Fig. S9.**
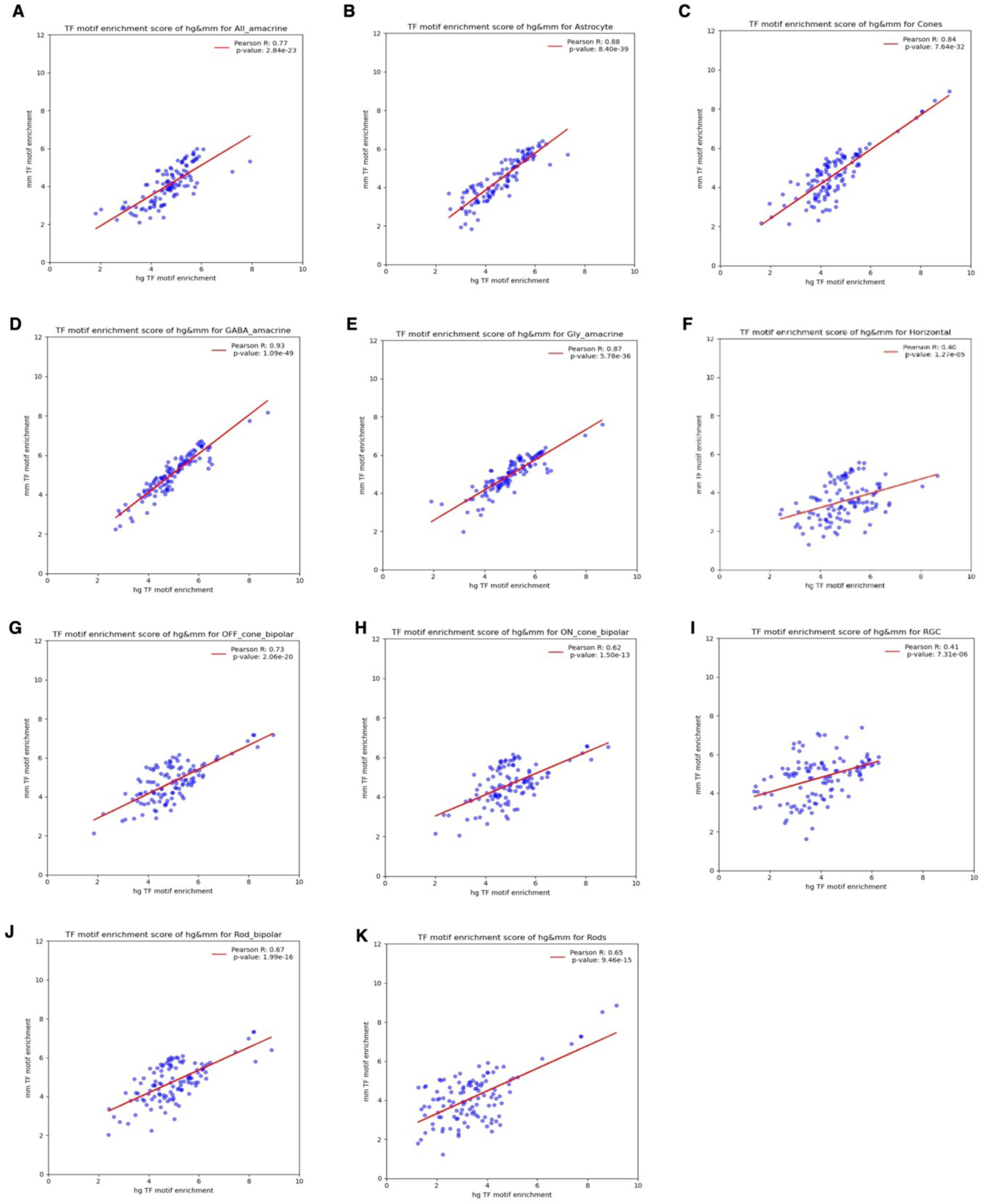
The conserved TF motifs of human and mouse retina for each cell type and the PCC of the enrichment score. (**A**) All amacrine **(B)** Astrocyte (**C)** Cones **(D)** GABA amacrine **(E)** Gly amacrine **(F)** Horizontal **(G)** OFF cone bipolar **(H)** ON cone bipolar **(I)** RGC **(J)** Rod bipolar **(K)** Rods.

**Fig. S10.**
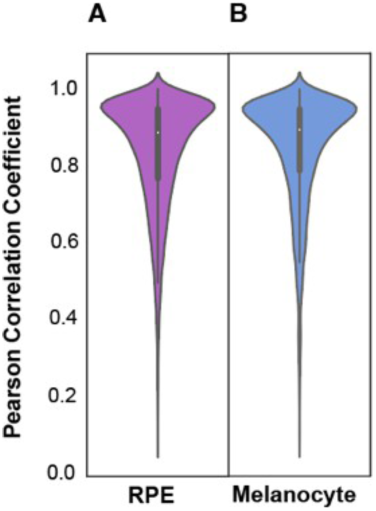
The violin plot of Pearson Correlation Coefficient between deep learning model predicted open chromatin in RPE and melanocytes and psuedobulk ATAC-seq data for each cell type on its validation dataset. (**A**) Pseudo bulk RPE **(B)** Pseudo bulk Melanocyte.

